# The biosynthesis, degradation, and function of cell wall β-xylosylated xyloglucan mirrors that of arabinoxyloglucan

**DOI:** 10.1101/2023.03.03.528403

**Authors:** L. F. L. Wilson, S. Neun, L. Yu, T. Tryfona, K. Stott, F. Hollfelder, P. Dupree

## Abstract

- Xyloglucan is an abundant polysaccharide in many primary cell walls and in the human diet. Decoration of its α-xylosyl side chains with further sugars is critical for plant growth, even though the sugars themselves vary considerably between species. Plants in the Ericales order—prevalent in human diets—exhibit β1,2-linked xylosyl decorations. The biosynthetic enzymes responsible for adding these xylosyl decorations, as well as the hydrolases that remove them in the human gut, are unidentified.
- GT47 xyloglucan glycosyltransferase candidates were expressed in Arabidopsis and *endo*-xyloglucanase products from transgenic wall material were analysed by electrophoresis, mass spectrometry, and NMR. The activities of gut bacterial hydrolases *Bo*GH43A and *Bo*GH43B on synthetic glycosides and xyloglucan oligosaccharides were measured by colorimetry and electrophoresis.
- *Cc*XBT1 is a xyloglucan β-xylosyltransferase from coffee that can modify Arabidopsis xyloglucan and restore the growth of galactosyltransferase mutants. Related *Vm*XST1 is a weakly active xyloglucan α-arabinofuranosyltransferase from cranberry. *Bo*GH43A hydrolyses both α-arabinofuranosylated and β-xylosylated oligosaccharides.
- *Cc*XBT1’s presence in coffee and *Bo*GH43A’s promiscuity suggest that β-xylosylated xyloglucan is not only more widespread than thought, but might also nourish beneficial gut bacteria. The evolutionary instability of transferase specificity and lack of hydrolase specificity hint that, to enzymes, xylosides and arabinofuranosides are closely resemblant.

## Introduction

All plant cells possess a primary cell wall (Kumar & Turner, 2015). In eudicots, xyloglucan (a hemicellulosic polysaccharide) makes up around 20–25 % of the primary cell wall dry weight, and is thought to play roles in cell wall integrity and extensibility, most likely through nuanced interactions with cellulose microfibrils (Hsieh *et al*., 2009; Cosgrove, 2022). Like cellulose, xyloglucan has a backbone of β1,4-linked glucosyl residues. However, unlike cellulose, the majority of these backbone residues are decorated with α1,6-linked xylose. These decorations typically appear in repeating patterns along the backbone—in most eudicots, for instance, every fourth glucose is left undecorated (Schultink *et al*., 2014). These xylosyl residues are frequently decorated with further sugar substituents at the C2 hydroxyl, as shown in Fig. 1a (along with their shorthand nomenclature (Fry *et al*., 1993), which is used below). The xylosyl residue may be substituted at C2 with β-galactose (β-Gal), β-galacturonic acid (β-GalA), α-arabinofuranose (α-Ara*f*), α-arabinopyranose (α-Ara*p*), or β-xylose (β-Xyl) to produce the ‘L’, ‘Y’, ‘S’, ‘D’, or ‘U’ structures, respectively (Fig. 1a). Furthermore, fucosylation of the terminal galactosyl, galacturonosyl or arabinopyranosyl residues produces the ‘F’, ‘Z’, and ‘E’ structures (Schultink *et al*., 2014).

**Figure 1.**
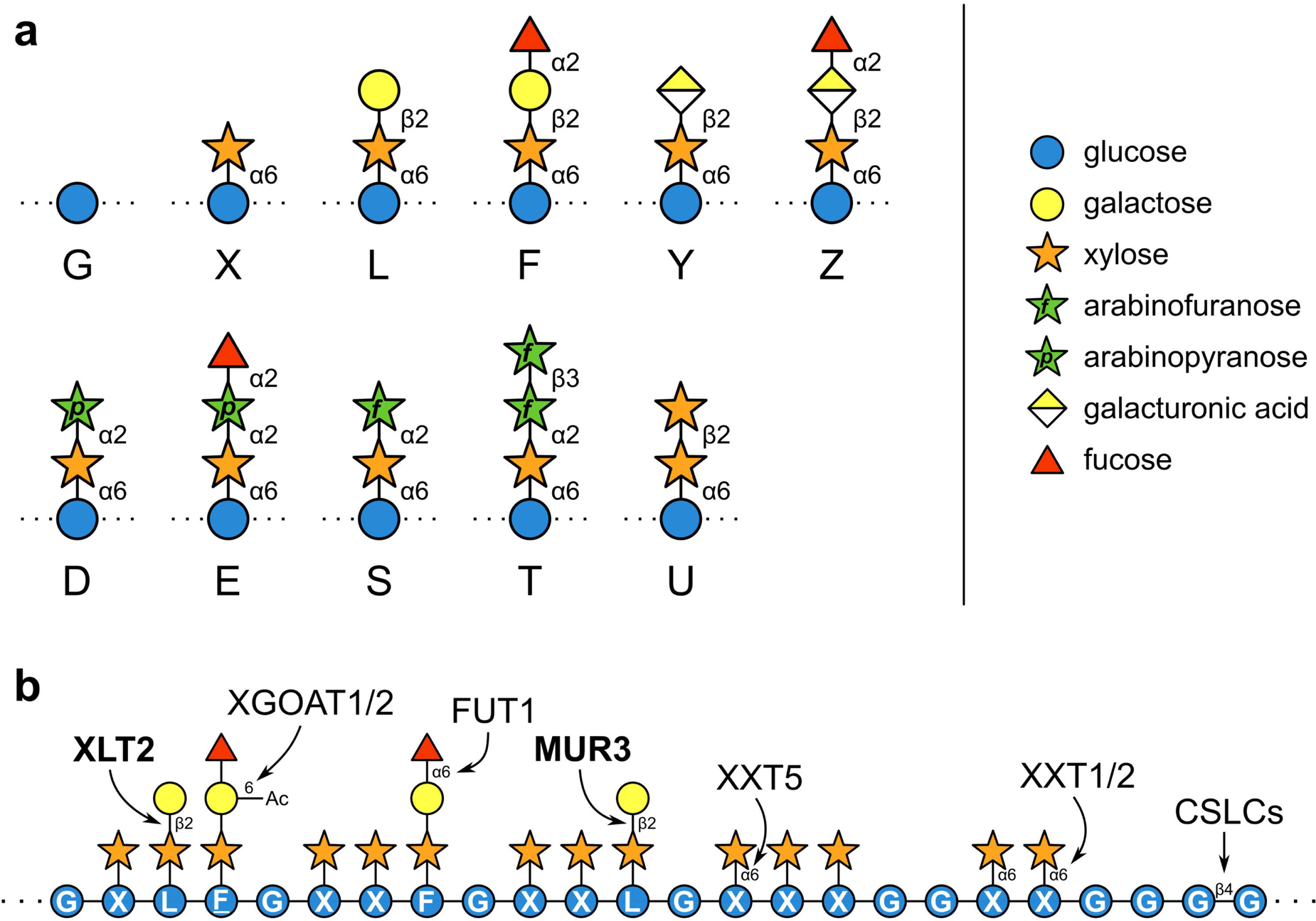
Xyloglucan sidechain nomenclature and biosynthesis. **a** To describe xyloglucan structures in shorthand, a letter is assigned for every backbone glucosyl residue according to its side chain (Fry *et al*., 1993). For instance, unsubstituted glucose is represented by ‘G’, whereas glucose substituted with a single xylosyl residue is termed ‘X’. **b** Enzymes involved in synthesising typical (fucogalacto)xyloglucan in the Arabidopsis Golgi body. Members of the GT47-A family are highlighted in bold.

In Arabidopsis and many other eudicots, the xyloglucan produced by most tissues appears to be predominantly composed of units of XXXG, XLXG, XXFG, and XLFG (Zabotina, 2012) (Fig. 1b). More generally, however, plants exhibit astonishing diversity in xyloglucan structure across the kingdom—even between different tissues. The large group of eudicots known as asterids, for example, display more unusual structures. As in monocots, some asterids exhibit xyloglucan based on an XXGG or XXG*_n_* repeat, as opposed to XXXG (Hoffman *et al*., 2005; Hsieh & Harris, 2009). Furthermore, asterid xyloglucans often contain dipentosyl side chains: for instance, the S structure (Ara*f*-α1,2-Xyl-α1,6-Glc) has been found in olive (Lamiales), oleander (Gentianales), and many species from the Solanales order, whereas the U structure of β-xylosylated xyloglucan (Xyl*p*-β1,2-Xyl-α1,6-Glc) has so far been found in bilberry and argan (Ericales) (Schultink *et al*., 2014). Dixylosyl side chains have also been reported in tobacco and aubergine (Solanales), but, at least in the latter, the terminal xylose was proposed to be α-linked based on its insensitivity to a β-xylosidase (Sims *et al*., 1996; Kato *et al*., 2010). The D structure (Ara*p*-α1,2-Xyl-α1,6-Glc) and its derivatives, on the other hand, have only been detected in non-spermatophytes (Zavyalov *et al*., 2019).

Xyloglucan is synthesised by glycosyltransferases (GTs) and acetyltransferases in the Golgi apparatus before being further modified by glycosidases and transglycosylases in the cell wall (Pauly & Keegstra, 2016). The biosynthesis of Arabidopsis (fucogalacto)xyloglucan is particularly well characterised: the glucan backbone is made by CSLC4,5,6,8 or 12 (CAZy family GT2), backbone glucosyl residues are xylosylated by XXT1–5 (GT34), xylosyl residues are galactosylated by XLT2 and MUR3 (GT47 subclade A), and galactosyl residues are fucosylated by FUT1 (GT37) and acetylated by XGOAT1/2 (Fig. 1b) (Campbell *et al*., 1997; Coutinho *et al*., 2003; Pauly & Keegstra, 2016; Zhong *et al*., 2018; Kim *et al*., 2020; Drula *et al*., 2022; Julian & Zabotina, 2022). The polysaccharide is then trimmed and remodelled in the cell wall by enzymes such as XTH endohydrolases/transglycosylases, AXY8 α-fucosidase, BGAL10 β-galactosidase, and XYL1 α-xylosidase (Pauly & Keegstra, 2016). However, this model cannot be easily extrapolated to other species, as, despite a close evolutionary relationship, homologues of the XLT2 and MUR3 galactosyltransferases in family GT47-A exhibit a remarkable variability in substrate specificity, both between paralogues and between orthologues from across the plant kingdom. For instance, two XLT2 orthologues in tomato, XST1 and XST2, decorate xylosyl residues with arabinofuranose, rather than galactose, to create the S structure (Schultink *et al*., 2013), whereas a paralogue in *Physcomitrium patens*, XDT, instead transfers arabinopyranose, creating the D structure (Zhu *et al*., 2018). Additionally, the hypothetical β-xylosyltransferase needed to synthesise the U structure has been suggested to belong to this family (Schultink *et al*., 2014). Even in Arabidopsis, an XLT2/MUR3 homologue expressed in root hair cells decorates the xylosyl residues with galacturonic acid, resulting in the Y side chain (Peña *et al*., 2012). Furthermore, some enzymes in the GT47-A family specifically recognise different polysaccharide acceptors: MBGT1 decorates galactoglucomannan with β-galactose for instance (Yu *et al*., 2022). Currently, there is no way to predict the activities of these enzymes from amino acid sequence, and current phylogenetic classifications fail to delineate the different donor substrate specificities. Without more functional characterisation, the inability to assign a substrate specificity from sequence prevents us from predicting the carbohydrate structures present in any given plant. Furthermore, we do not yet know whether the rapid evolution of GT47-A activity is purely a product of inconsequential drift (perhaps resulting from a weakly defined substrate specificity that is easily perturbed by active site mutations, or the promiscuity of a recent common ancestor) or whether it is driven by environmental selection for wall polysaccharides with altered physical or biotic attributes.

It is plausible that such structural diversity could provide resistance against cell wall-degrading pathogens (Malinovsky *et al*., 2014) and constitute an evolutionary selection criterion. The xyloglucan degradation system of *Xanthomonas citri* pv. *citri* 306 (Vieira *et al*., 2021) is currently the best characterised pathogen model for xyloglucan utilisation. Interestingly, this gene cluster appears only to encode enzymes for degradation of fucogalacto- and acetoxyloglucan (though two co-regulated GH43-family hydrolases exhibit activity on synthetic α-arabinofuranosides). Hence, it remains to be determined whether the specialised plant pathogens in this genus are able to metabolise other types of xyloglucan, such as the β-xylosylated xyloglucan of plants in the Ericales order.

With xyloglucan making up perhaps as much as 20% of the dry weight of some fruit and vegetables (Toushik *et al*., 2017), the same structural complexity ought to have consequences for the fibre-degrading microbiota in our guts (and therefore indirectly for human health) (Koropatkin *et al*., 2012). Interestingly, the ability to degrade xyloglucan appears to be a rare trait amongst gut micro-organisms (Hartemink *et al*., 1996; Larsbrink *et al*., 2014). The best-studied gut xyloglucan degradation system is that encoded by the xyloglucan utilisation locus (XyGUL) of *Bacteroides ovatus* ATCC 8483 (Larsbrink *et al*., 2014). The *B. ovatus* XyGUL encodes both secretory *endo*-xyloglucanases, which cleave extracellular xyloglucan into xyloglucan oligosaccharides (XyGOs), and *exo*-glycosidases, which disassemble XyGOs after their import into the periplasm. Amongst the *exo*-glycosidases are two GH43 α-arabinofuranosidases: *Bo*GH43A and *Bo*GH43B. Importantly, these are the only enzymes demonstrated to exhibit α-arabinofuranosidase activity on xyloglucan or XyGOs. Although the *B. ovatus* XyGUL does not encode an α-fucosidase, other Bacteroidetes XyGULs encode functional GH95 α-fucosidases, leading to the suggestion that individual XyGULs are tailored to specific types of xyloglucan (Larsbrink *et al*., 2014; Déjean *et al*., 2019). However, it is currently unknown whether gut micro-organisms are capable of metabolising xyloglucans other than fucogalacto- and arabinoxyloglucan.

The diversity of xyloglucan structures across the plant kingdom has yet to be fully explored. At the same time, the range of activities exhibited by GT47-A family members has not been completely characterised. Furthermore, despite the widespread interest in the role of dietary polysaccharides such as xyloglucan in gut health, we lack comprehensive knowledge of their metabolic fate. These factors limit not only our ability to understand xyloglucan function and versatility *in planta* but also to predict, select, and design xyloglucans with beneficial material and health properties. Here, making use of molecular phylogeny, recently released AlphaFold structures (Jumper *et al*., 2021; Varadi *et al*., 2022), and functional characterisation *in planta*, we explore the evolution and diversity of donor sugar specificities in the GT47-A family. Accordingly, we identify a xyloglucan β-xylosyltransferase (*Cc*XBT1) from *Coffea canephora* (Robusta coffee, from outside of the Ericales order) that is closely related to previously characterised xyloglucan α-arabinofuranosyltransferases. We use *Cc*XBT1 to make a novel xyloglucan in Arabidopsis composed almost entirely out of XUXG units that can functionally replace native fucogalactoxyloglucan. To demonstrate this, we present a glycosidase-based system for analysing xyloglucan structure by electrophoresis. Furthermore, we use the novel xyloglucan structure to investigate the activity of *Bo*GH43A. We find that *Bo*GH43A, and perhaps *Bo*GH43B, are promiscuous and therefore likely act as dual-purpose α-arabinosidases/β-xylosidases in *B. ovatus* xyloglucan metabolism. Hence, both the synthesis and degradation of β-xylosylated xyloglucan are closely related with those of arabinoxyloglucan, suggesting that, in general, the evolutionary leap between activity on L-arabinosides and activity on D-xylosides requires relatively few mutations, therefore constituting a shortcut in the landscape of enzyme evolution.

## Materials and methods

### Bioinformatics

GT47-A sequences were obtained from available genomic datasets (Table S1) using HMMER (Eddy, 2011) as previously described (Yu *et al*., 2022) with an *E* value threshold of 1×10^−70^. Where only nucleotide sequences were available, we searched for GT47-A sequences with TBLASTN (Altschul *et al*., 1990), using all the Arabidopsis GT47-A protein sequences as individual queries; redundant hits were removed by CD-HIT clustering (Li & Godzik, 2006; Fu *et al*., 2012) with a 95% similarity threshold. TBLASTN hits were named *Xx*GT47-A_1, *Xx*GT47-A_2 etc.. All sequences were then truncated to the GT47 domain (residues 123–512 in *At*XLT2) as previously described (Wilson *et al*., 2022). Following this, sequences shorter than that of *At*XLT2 by more than 33% were removed. Model selection and phylogenetic inference were then achieved using IQ-TREE (Nguyen *et al*., 2015) with 1,000 ultra-fast bootstrap pseudo-replicates (Hoang *et al*., 2018). Subtrees were extracted using FigTree (Rambaut & Drummond, 2018). Columns corresponding to the fourteen residues of interest in *At*XLT2 were extracted, concatenated, and submitted to the WebLogo 3 (Crooks *et al*., 2004) server (http://weblogo.threeplusone.com/).

### Molecular biology

The coding sequences of *Vm*XST1, *Cc*XBT1 (Cc07_g06550), and Cc07_g06570 (Table S2) were synthesised *de novo* by GENEWIZ (Suzhou, China) as Golden Gate MoClo/Phytobrick B3–B4 CDS parts (Patron *et al*., 2015). The promoters of XXT2 and CESA3 and the coding sequence of *Sl*XST1 were amplified from Arabidopsis or tomato genomic DNA, respectively, by PCR using Q5 High-Fidelity DNA Polymerase (New England Biolabs, Ipswich, MA, USA; primers are listed in Table S3). Constructs and transgenic Arabidopsis lines were then generated as previously described (Lyczakowski *et al*., 2021).

Plasmids for expression of *Bo*GH43A (pET-YSBLIC-*Bo*GH43A) and *Bo*GH43B (pET21a-BoGH43B) were a gift of Gideon Davies (York, UK). Protein expression and purification was performed according to Hemsworth et al. (2016), with minor modifications (Methods S1).

### Plant growth, mutants, and sourced material

All plants were Col-0 in ecotype. The *mur3-1* (CS8566), *mur3-3* (SALK_141953), and *xlt2* mutants (GABI_552C10) were obtained from the Nottingham Arabidopsis Stock Centre. Plants were grown on a 9:1 mix of Levington M3 compost and vermiculite and grown at 21°C under a 16 h / 8 h photoperiod.

Crosses to produce the *xlt2 mur3-1* double mutant were achieved by the method of Weigel & Glazebrook (2006), with genotyping as described by Jensen *et al*. (2012) but using the primers 5’-GCATCTACCCAAGAACTACACAACCGACT-3’ and 5’-GTCGCTCCTCGATGCTTATGTTTC-3’ and the restriction enzyme HinfI.

Leaves of olive (*Olea europaea*), argan (*Argania spinosa* / *Sideroxylon spinosum*), Arabica coffee (*Coffea arabica* ‘Catimor’), *Catharanthus roseus*, and borage (*Borago officinalis*) were a gift of the Cambridge University Botanic Garden. Blueberry and kiwi fruits were purchased from a supermarket. Green cherry tomato fruit (‘Tumbling Tom Red’) was a gift of Mrs Elaine Lundy Wilson (Canterbury, UK).

### Xyloglucan preparation and digestion

De-acetylated hemicelluloses were extracted from alcohol insoluble residue (AIR) as described previously (Goubet *et al*., 2009; Yu *et al*., 2022). For *endo*-xyloglucanase digestions, 2 μl 14 μM xyloglucan-specific *endo*-β1,4-glucanase (XEG) from *Aspergillus aculeatus* (Pauly *et al*., 1999) (*Aa*XEG; Novozymes, Copenhagen, Denmark) was added to 70 μl PD-10 eluate (equivalent to 0.6 mg AIR) and incubated at 37 °C for 18 h (in the case of *C. arabica* leaves, 16 μl *Aa*XEG was added to 187.5 μl PD-10 eluate). Digests were dried using a centrifugal evaporator before resuspending oligosaccharides in 200 μl 65 % ethanol and incubating at −20 °C overnight. The samples were centrifuged at 10,000 × *g* for 15 min before drying the supernatant and resuspending the oligosaccharides in 1 ml 50 mM ammonium acetate, pH 6.0.

*Exo*-glycosidases (Table S4) were added directly to 100 μl resuspended oligosaccharides (PACE implied that XyGO concentration was in the low micromolar range). Reactions were carried out at 37 °C for 18 h, except in the case of *Cg*GH3 digestions, which required 48 h for completion. Since we found *Cg*GH3 to exhibit a secondary β-galactosidase activity, this enzyme could not be used to analyse XyGOs that had not already been treated with Fam35 β-galactosidase. In sequential digestions, α-xylosidase was deactivated by ethanol precipitation (as described above), whereas other enzymes were removed by passing the sample through a 3 kDa NanoSep centrifugal filter device (VWR, Radnor, PA, USA).

### PACE

Polysaccharide analysis by carbohydrate electrophoresis (PACE) was carried out as described by Goubet et al. (2002). One hundred microlitres of oligosaccharide mixture was used for each sample, along with 5 μl labelling reagent and 10 μl 6 M urea for final resuspension. The 1,000-V step running step was extended to 150 min. Cello-oligosaccharide standards (50 pmol each) were obtained from Megazyme (Bray, Ireland).

### Size exclusion chromatography of oligosaccharides

Size-exclusion chromatography was carried out by a similar method to Yu *et al*. (2022) in 50 mM ammonium acetate, pH 6.0 (Methods S2). Eluates required clean-up with a PD MiniTrap G-10 column (Cytiva, Marlborough, MA, USA).

### Spectroscopy

Dried XyGOs (without further desalting), or reductively aminated oligosaccharides (passed through a GlycoClean S column) were analysed by mass spectrometry (MS) and tandem MS–MS, respectively, as previously described (Tryfona & Stephens, 2010; Yu *et al*., 2022; Wilson *et al*., 2022). Data were acquired on reflector positive ion mode (mass range 300– 2,000 Da). Solution-state NMR was carried out as previously (Yu *et al*., 2022), with the spectrometer operating at 800 MHz.

### Monosaccharide analysis

Following NMR, approximately 25 μg of the purified XUXG oligosaccharide was dried, resuspended in 400 μl 2 M trifluoroacetic acid (TFA), and incubated at 120 °C for 1 h. After drying, the sample was resuspended in 200 μl MilliQ water and injected into a Dionex ICS300 HPLC system alongside 10–400 μM monosaccharide standards. High-performance anion-exchange chromatography with pulsed amperometric detection (HPAEC-PAD) then proceeded as described previously (Bromley *et al*., 2013).

### Kinetic enzyme assays

Initial substrate screens on *p*NP-α-Ara*f* (Merck), *p*NP-β-Xyl (Merck), and *p*NP-α-Ara*p* (Glycosynth, Warrington, UK) (Figs. 5c and S8) were performed in 10 mM HEPES, pH 7, 250 mM NaCl using 1 μM enzyme and 1 mM *p*NP-glycoside (4-nitrophenyl-glycoside), measuring the product absorbance at A_405_ every 5 s for 5 h and then once again after 19 h in a plate reader. To determine pH optima, the activities of *Bo*GH43A and *Bo*GH43B were measured over a range of pH conditions. The following buffers were prepared at 100 mM concentration with pH 0.5 increments between the given ranges: citrate-phosphate (pH 3.0– 7.0), HEPES (pH 7.0–8.0), Tris-HCl (pH 8.0–9.0), glycine (pH 9.0–10.5). Initial reaction velocities were recorded by measuring A_405_ with 1 mM substrate (*p*NP-β-Xyl for *Bo*GH43A and *p*NP-α-Ara*f* for *Bo*GH43B, respectively) and 2 μM enzyme in 50 mM buffer. Activities were normalised to the highest reaction velocity measured for each enzyme and plotted against the pH. Detailed kinetic experiments to obtain Michaelis-Menten parameters for *p*NP-α-Ara*f* and *p*NP-β-Xyl (Fig. S9 and Table S6) were performed in 50 mM HEPES, pH 7.5, with 1 μM enzyme at 20 °C. Reaction progress was followed by measuring A_405_ every 1 min in a plate reader to measure initial rates. Controls for non-enzymatic hydrolysis were carried out in the absence of enzyme and the background rates subtracted from the initial rates measured for the enzymatic reactions.

## Results

### The GT47-A_III_ clade is expanded in asterid genomes and exhibits subgroups with altered putative sugar-binding residues

Glycosyltransferases from GT47-A exhibit an unusually wide range of substrate specificities, with the sugar nucleotides UDP-Gal, UDP-GalA, UDP-Ara*f*, and UDP-Ara*p* all known to be employed as donor substrates by different members. This variability likely underlies the variety of xyloglucan side chain structures observed in the cell walls of asterids (Schultink *et al*., 2014; Pauly & Keegstra, 2016). Within asterid GT47-A enzymes specifically, activities have already been demonstrated for tomato galactosyltransferase *Sl*MUR3 (from group GT47-A_VI_, as classified by Yu *et al*., 2022) and arabinofuranosyltransferases *Sl*XST1 and *Sl*XST2 (GT47-A_III_) (Schultink *et al*., 2013); however, the enzymatic basis for asterid xyloglucan structural diversity is yet to be fully explored. To investigate whether asterids might possess an enlarged complement of GT47-A enzymes, we calculated a phylogeny of asterid GT47-A protein sequences, with a focus on Ericales and lamiids (Fig. S1). Interestingly, in line with previous, smaller-scale phylogenies (Schultink *et al*., 2013), the asterid proteomes were observed to exhibit a substantial expansion in group GT47-A_III_ (the clade containing *At*XLT2, *Os*XLT2 and *Sl*XST1/2) relative to the non-asterid references. This expansion hints at potential neofunctionalisation within the clade.

To investigate further, we examined the GT47-A_III_ subtree in more detail. Three distinct subclades were plainly visible within the asterid sequences; we named these subclades *a*–*c* (GT47-A_III_*a*–*c*; Figs. 2a, S1, and S2). *At*XLT2 was apparently grouped in subclade *a*, whereas subclades *b* and *c* were specific to asterids. Subclade *b* contained the tomato sequence Solyc02g092840, which was previously expressed in Arabidopsis but not found to exhibit an obvious activity (Schultink *et al*., 2013). Within the lamiid sequences, subclade *c* was clearly split into two lower-level subclades (*c*1 and *c*2). Although GT47-A_III_*c*1 contains both *Sl*XST1 and *Sl*XST2, no tomato sequences were grouped into GT47-A_III_*c*2.

**Figure 2.**
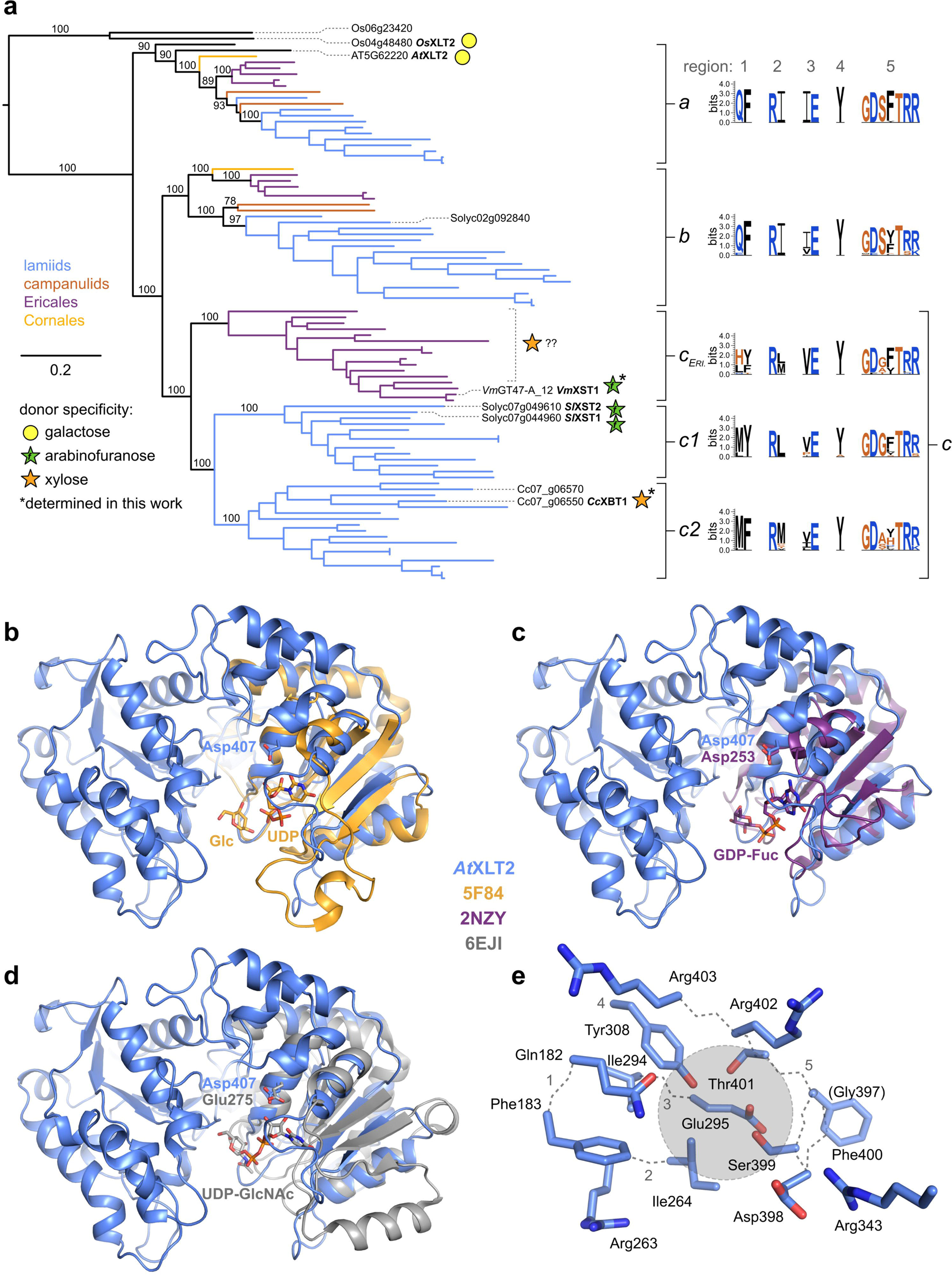
Despite their close phylogenetic grouping, members of GT47-A_III_ exhibit an unexpected variability in activity that might stem from subtle changes in three regions of interest. **a** Maximum-likelihood phylogeny of GT47-A subclade III (GT47-A_III_; XLT2-related) sequences from a range of asterid species, as well as the following non-asterids references: Arabidopsis, rice, and *Aquilegia coerulia*. Known donor substrate specificities are annotated with sugar symbols (as designated in the key). Ericales GT47-A_III_*c* (*c_ERI._*) sequences are loosely and tentatively labelled with xylose because the glycosyltransferase responsible for β-xylosylating xyloglucan in argan, bilberry, and blueberry has not yet been identified but is likely to belong to this clade. Sequences were truncated to the GT47 domain prior to phylogenetic inference using IQ-TREE. Horizontal branch lengths indicate average number of subsititutions per site (refer to scale bar). Support values at important splits represent percentage replication within 1,000 ultra-fast bootstrap pseudo-replicates. The tree shown is a subtree of the full GT47-A phylogeny (Fig. S1). See Fig. S2 for taxon labels and further support values. For sites corresponding to putative donor-sugar-binding residues in *At*XLT2, the conservation was examined by plotting a sequence logo for each phylogenetic subgroup using WebLogo 3. **b–d** AlphaFold model of *At*XLT2 (blue) aligned by its C-terminal subdomain to the three most similar nucleotide-sugar-bound GT-B structures according to the DALI webserver (http://ekhidna2.biocenter.helsinki.fi/dali/): **b** UDP and glucose bound to Drosophila POGLUT1 (PDB: 5F84; GT90), **c** GDP-fucose bound to *Helicobacter pylori* FucT (PDB: 2NZY; GT10), **d** UDP-*N*-acetylglucosamine bound to *Campylobacter jejuni* PglH (PDB: 6EJI; GT4). The conserved ribose-binding aspartate/glutamate residue is indicated where present. For clarity, only the C-terminal subdomain and bound nucleotide sugar is shown for each of the aligned structures. **e** Close-up of putative *At*XLT2 donor sugar binding pocket. Grey circle represents approximate position of donor sugar in aligned structures. Grey dotted lines trace the main chain backbone and are numbered according to the five regions of interest.

To screen for potential new donor sugar specificities within these groups, we sought to identify likely substrate-binding residues, using the AlphaFold-predicted structure of *At*XLT2 as a model. From DALI searches (http://ekhidna2.biocenter.helsinki.fi/dali/), the most similar experimentally determined sugar-nucleotide-bound structures were found to be from families GT90, GT10, and GT4, and superimposing them on the *At*XLT2 model consistently aligned their ligands in a conserved pocket (Fig. 2b–e). In each case, the aligned nucleotide was placed below the side chain of Asp407—a highly conserved residue that has since been shown to bind sugar nucleotides in other GT47 family members (Leisico *et al*., 2022; Wilson *et al*., 2022; Li *et al*., 2023); hence, this is the likely binding pocket for UDP-Gal in *At*XLT2.

Following this argument, we focused on residues in *At*XLT2 within 5 Å of any of the three aligned donor sugars, which comprised five different regions of interest (Arg343 was omitted for steric reasons). Together, they formed a QF-RI-IE-Y-GDSFTRR pattern, which was fully conserved in *Os*XLT2 and almost fully conserved in GT47-A_VI_ members *At*MUR3, *Sl*MUR3, and *Os*MUR3 (which exhibited a QF-RI-**V**E-Y-GDS**Y**TRR pattern). Hence, conservation of these residues may be important for the shared galactosyltransferase activity (Schultink *et al*., 2013; Liu *et al*., 2015) of all five enzymes.

To investigate the variability of these residues within the different subclades of GT47-A_III_, we created a sequence logo for each subclade. Interestingly, while the general pattern seen in *At*XLT2 and other characterised galactosyltransferases was strongly conserved in subclades *a* and *b*, subclade *c* exhibited noticeable differences, not only with respect to subclades *a* and *b* but also between its lower-level subclades (Fig. 2a). Notably, and consistent with our hypothesis, the arabinofuranosyltransferases *Sl*XST1 and *Sl*XST2 exhibited patterns of **MY**-R**L**-**V**E-Y-GD**G**FTRR and **MY**-R**L**-**V**E-Y-GD**GL**TRR, respectively. Hence, we predicted that family members with unusual substitutions in these regions could potentially harbour novel activities.

### GT47-A_III_c enzymes CcXBT1 and VmXST1 are xyloglucan pentosyltransferases capable of rescuing xyloglucan galactosyltransferase mutant phenotypes

To explore the range of enzyme activities in GT47-A_III_, we selected three sequences for further characterisation based on their putative donor-sugar-binding residues: (1) Cc07_g06550 (GT47-A_III_*c*2; **L**F-R**S**-IE-Y-GDS**V**TRR) from robusta coffee (*C. canephora*), hereafter *C. canephora* xyloglucan β-xylosyltransferase 1 (*Cc*XBT1); (2) Cc07_g06570 (GT47-A_III_*c*2; **M**F-R**V**-**L**E-Y-GDSFTRR), also from *C. canephora*; and (3) *Vm*GT47-A_12 (GT47-A_III_*c*; **LY**-R**M**-**V**E-Y-GD**G**FTRR) from cranberry (*Vaccinium macrocarpon*), hereafter *Vm*XST1. To screen for potential xyloglucan-modifying activity, we tested the capacity of each enzyme to rescue the phenotype of the Arabidopsis *mur3-3* mutant, which lacks MUR3 activity and exhibits a dwarfed, cabbage-like growth phenotype (Madson *et al*., 2003; Kong *et al*., 2015). *Cc*XBT1, Cc07_g06570, and *Vm*XST1 were each expressed under the promoter of XXT2 (one of the main xyloglucan α-xylosyltransferases in Arabidopsis (Julian & Zabotina, 2022)). We deleted a stretch of repetitive and putatively disordered residues (residues 167–193) from the *Vm*XST1 stem domain in order to maximise chances of expression. After six weeks of growth, we compared the growth phenotypes of T_1_ *mur3-3* transgenic plants to wild-type and untransformed *mur3-3* plants. Although expression of Cc07_g06570 did not appear to alter the phenotype, expression of *Cc*XBT1 rescued growth almost to wild-type levels (Fig. S4). Expression of *Vm*XST1, however, had an intermediate effect on growth, which was variable between different lines. These results are consistent with the idea that *Cc*XBT1 and potentially *Vm*XST1 are able to alter xyloglucan structure.

To better characterise the products of *Cc*XBT1 and *Vm*XST1, we wanted to express these enzymes in a background lacking galactosylation from both MUR3 and XLT2. Because of the prohibitively strong phenotype of *mur3-3*, we crossed *xlt2* plants with a weaker allele, *mur3-1*, whose growth is less severely stunted (*mur3-1* plants express a MUR3 point mutant with very low activity) (Madson *et al*., 2003; Tamura *et al*., 2005). The resultant *xlt2 mur3-1* double homozygous mutant has been previously characterised (Jensen *et al*., 2012; Kong *et al*., 2015) and used in many similar analyses (Schultink *et al*., 2013; Liu *et al*., 2015; Zhu *et al*., 2018). To maximise product yield, we expressed both *Cc*XBT1 and *Vm*XST1 under the strong, primary-cell-wall-synthesis-specific promoter of cellulose synthase 3 (CESA3). As a positive control, we made similar transgenic lines expressing *Sl*XST1. Once more, we compared the phenotypes of six-week-old T_1_ transgenic plants. Expression of either *Sl*XST1 or *Cc*XBT1 was sufficient to restore growth to wild-type levels (Figs. 3a and S5a). However, as before, expression of *Vm*XST1 resulted in only partial complementation. Nevertheless, these results are consistent with the idea that both enzymes modify xyloglucan.

**Figure 3.**
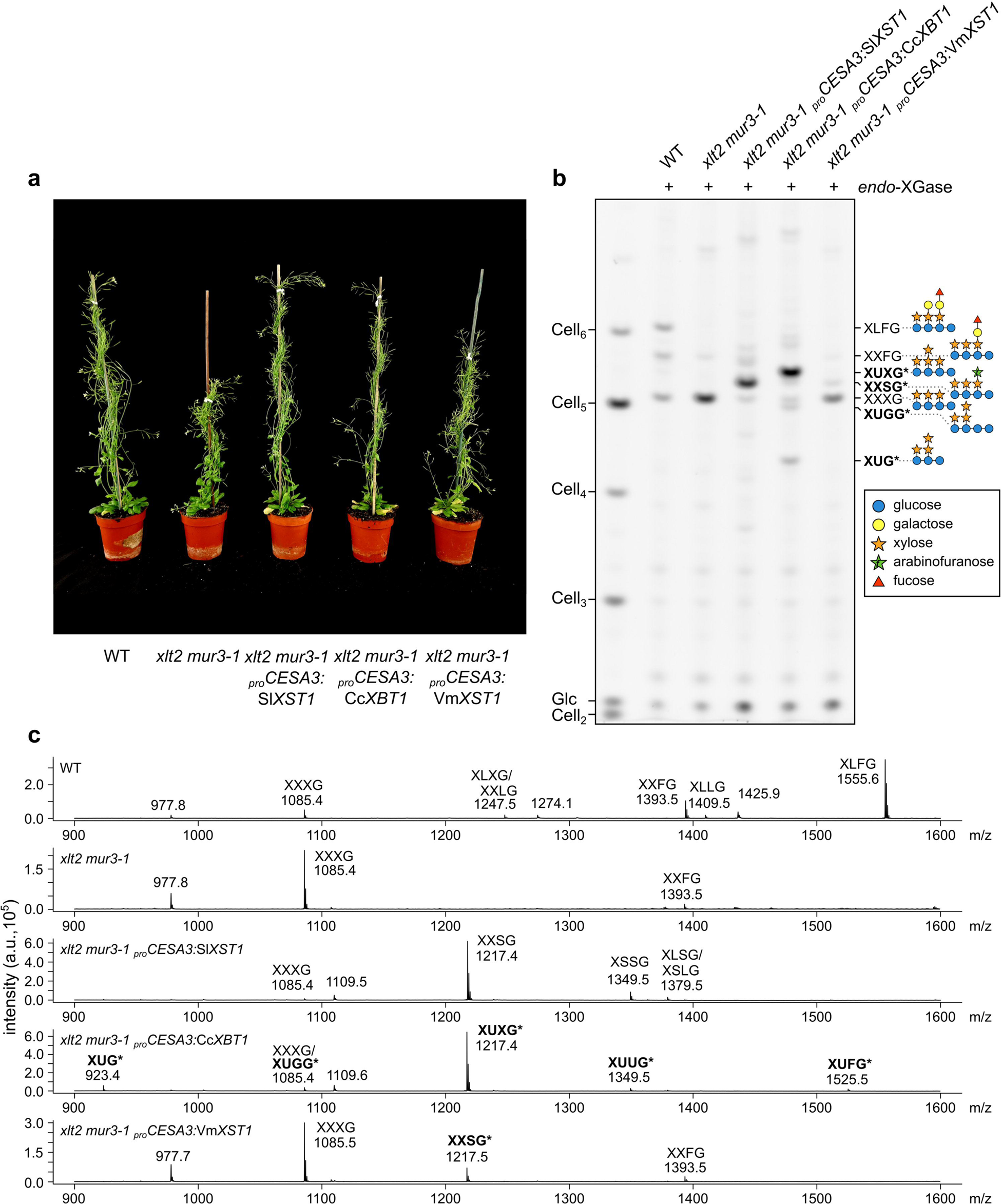
Expression of *Cc*XBT1 in *xlt2 mur3-1* Arabidopsis introduces a novel cell wall xyloglucan structure and rescues the mutant growth phenotype. **a** Representative six-week-old heterozygous transformants (T_1_ generation) expressing *Sl*XST1, *Cc*XBT1, or *Vm*XST1 under the strong primary wall-specific promoter of *CESA3*. Expression of *Sl*XST1 or *Cc*XBT1 fully rescues the mildly stunted phenotype of *xlt2 mur3-1*, whereas expression of *Vm*XST1 results in only partial complementation. See Fig. S5 for further transgenic lines. **b** Polysaccharide analysis by carbohydrate electrophoresis (PACE) analysis of *endo*-xyloglucanase (*endo*-XGase) products from transgenic cell wall material. Hemicellulose was extracted by alkali treatment of leaf alcohol insoluble residue (AIR) and digested with *Aa*XEG *endo*-XGase (which cleaves the xyloglucan backbone regularly at unsubstituted glucosyl residues). Products were subsequently derivatised with a fluorophore and separated by electrophoresis. Band assignments made later on in this work are marked with an asterisk. See Fig. S5 for further transgenic lines and Fig. S6 for no-enzyme controls. **c** MALDI-TOF mass spectrometry of the same *endo*-XGase products. Ions assigned in this work are marked with an asterisk. Note that, while PACE is fully quantitative, MALDI-TOF is only semi-quantitative.

To analyse the xyloglucan structures in the cell walls of these plants, we prepared alcohol-insoluble residue (AIR) from their rosette leaves and extracted hemicellulose using alkali. XyGOs were released by treatment with xyloglucan-specific *Aspergillus aculeatus endo*-β1,4-glucanase (*Aa*XEG; hereafter *endo*-XGase), which requires an unsubstituted backbone glucosyl residue at the −1 subsite (Pauly *et al*., 1999). The products were analysed by polysaccharide analysis by carbohydrate electrophoresis (PACE; Figs. 3b and S5–S6) and matrix-assisted laser desorption/ionisation–time of flight (MALDI-TOF) mass spectrometry (Fig. 3c). With reference to previous XyGO PACE band assignments (Yu *et al*., 2022), we determined that the predominant oligosaccharides released by *endo*-XGase from wild-type hemicellulose were XXXG, XXFG, and XLFG, as well as a smaller amount of XXLG and XLLG. In contrast, *xlt2 mur3-1* material released almost exclusively XXXG, as well as a very small amount of XXFG—as previously reported (Kong *et al*., 2015). However, compared to untransformed *xlt2 mur3-1*, *Sl*XST1- and *Cc*XBT1-expressing plants exhibited strikingly different XyGO profiles. The PACE profile from *Sl*XST1-expressing plants was dominated by a single band that corresponded to a MALDI-TOF ion with *m*/*z* 1,217—likely representing a (sodiated) octasaccharide composed of four pentoses and four hexoses. From the original characterisation of *Sl*XST1 in Arabidopsis, this XyGO is already known to be XXSG (Schultink *et al*., 2013), and interestingly, *Vm*XST1-expressing plants yielded a XyGO co-migrating with (and of identical mass to) the main *Sl*XST1 product (albeit of much lower abundance).

However, although the products released from *Cc*XBT1-expressing plants also exhibited a dominant band corresponding to the same mass, this main XyGO exhibited substantially different mobility in PACE analysis, indicating that it possesses a different structure. Furthermore, *Sl*XST1- and *Cc*XBT1-expressing plants differed in the presence/absence of several other bands and ions. Interestingly, two XyGOs from the *Cc*XBT1 material had higher mobility than XXXG in PACE analysis. Their smaller implied size indicates that they most likely originate from XXGG-type xyloglucan, which (when de-acetylated by alkali) can be cleaved on the reducing end side of either unsubstituted glucose residue by *endo*-XGase, producing XXG, XXGG, or decorated versions thereof.

### CcXBT1 is a β1,2-xylosyltransferase that transfers xylose to the second α-xylosyl residue in the XXXG repeat

To further characterise these potentially novel structures, we tested their sensitivity to two different *exo*-glycosidases. The first of these was *Cj*Abf51, an α-arabinofuranosidase whose primary activity is on α1,2- and α1,3-linked terminal arabinofuranosyl residues on xylan and arabinan (Beylot *et al*., 2001). Interestingly, both XXSG and the co-migrating oligosaccharide from *Vm*XST1-expressing plants could be converted to XXXG by this enzyme (Fig. 4a). Hence, we concluded that *Vm*XST1 is likely an arabinofuranosyltransferase with weak levels of activity, but the same specificity as *Sl*XST1 (at least when expressed in Arabidopsis). In contrast, the main XyGO from *Cc*XBT1-expressing plants was insensitive to *Cj*Abf51 treatment.

**Figure 4.**
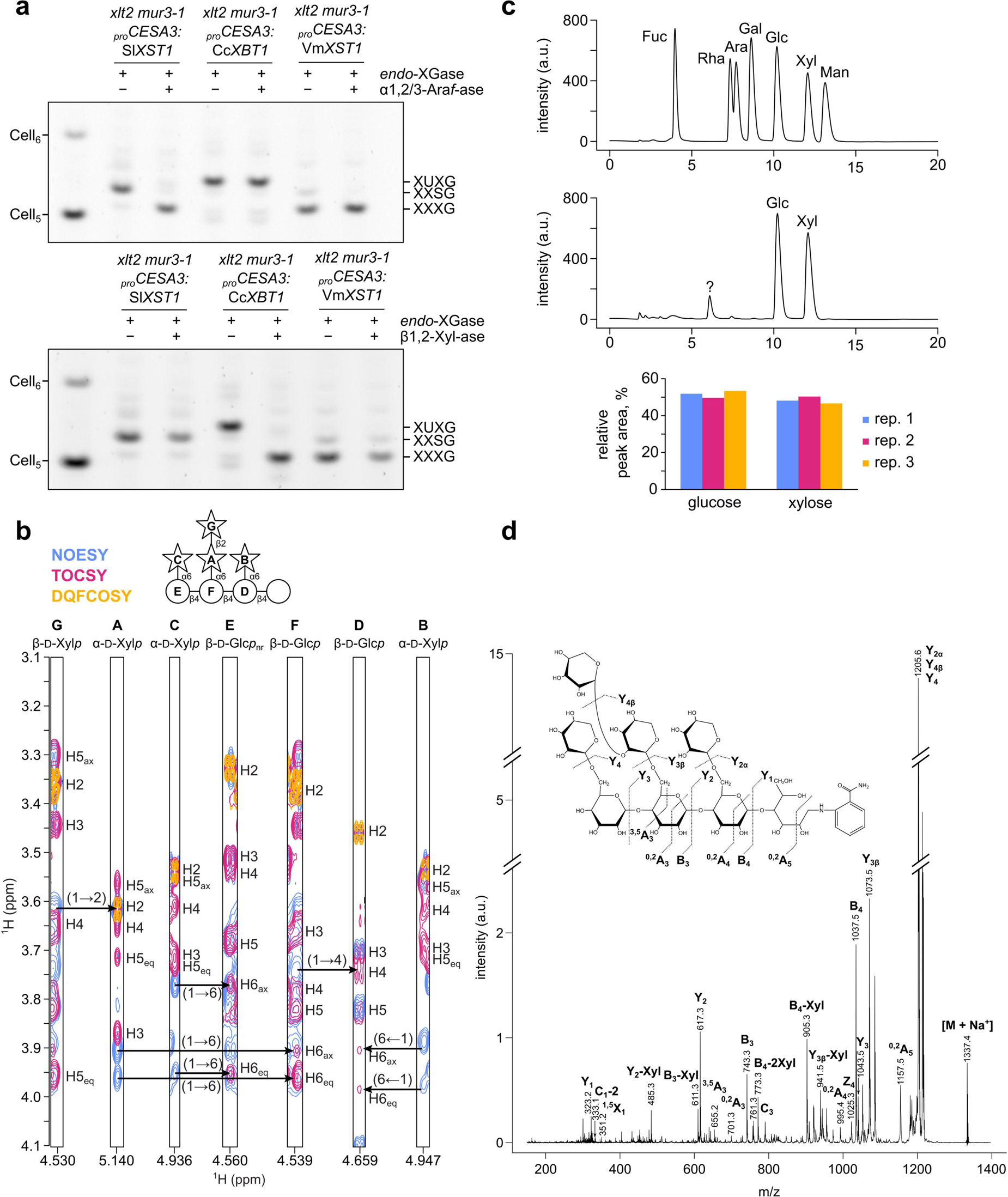
Characterisation of the XUXG endo-xyloglucanase product from *Cc*XBT1-expressing plants. **a** Sensitivity of endo-xyloglucanase products to α-arabinofuranosidase *vs* β-xylosidase (PACE gel). Alkali-extracted hemicellulose from transgenic plants was digested with *Aa*XEG *endo*-xyloglucanase. Following ethanol precipitation of undigested polymers and protein, the oligosaccharide products were treated with either *Cj*Abf51 α-arabinofuranosidase (α1,2/3-Ara*f*-ase) or *Cg*GH3 β-xylosidase (β1,2-Xyl-ase). Products were subsequently derivatised with a fluorophore and separated by electrophoresis. **b** Solution NMR analysis of purified XUXG oligosaccharide from *endo*-XGase digest of *Cc*XBT1-expressing plants. Each strip corresponds to a reference proton (C_1_–<u>H</u>) from one of seven distinct monosaccharides (**A**–**G**; stars = xylose, circles = glucose). The reducing end glucose was not visible in our analysis. Contours represent cross-peaks detected in ^1^H–^1^H NOESY (blue), TOCSY (pink), and DQFCOSY (yellow) experiments. Chemical shift assignments are provided in Table S5. **c** Monosaccharide analysis of the purified XUXG oligosaccharide following NMR. Monosaccharides were liberated using trifluoracetic acid (TFA) hydrolysis before separation and quantification with high-performance anion-exchange chromatography. Upper trace: 400 μM monosaccharide standards (representative of three technical replicates). Centre trace: monosaccharides released by TFA hydrolysis of the XUXG oligosaccharide;= unknown contaminant (representative of three technical replicates). Lower chart: relative quantification of glucose and xylose peak areas from XUXG hydrolysis (using standards as references). **d** Spectrum from tandem mass spectrometry (MS–MS) with collision-induced dissociation (CID) of purified XUXG oligosaccharide following reducing-end derivatisation with 2-aminobenzamide.

Secondly, we treated the XyGOs with *Cg*GH3, a β-xylosidase whose primary activity is on the β1,2-linked xylosyl residues of xylan D^2,3^ side chains (Xyl-β1,2-Ara*f*-α1,3-Xyl) (Tryfona *et al*., 2019). Conversely, whereas *Sl*XST1 and *Vm*XST1 XyGOs were insensitive to *Cg*GH3, the main *Cc*XBT1 XyGO could be completely converted to XXXG after extended treatment. This result suggested that *Cc*XBT1 may constitute a xyloglucan β-xylosyltransferase.

To confirm this hypothesis, we purified the main *Cc*XBT1 XyGO by size-exclusion chromatography and subjected it to two-dimensional solution NMR experiments, monosaccharide analysis, and collision-induced dissociation–mass spectrometry/mass spectrometry (CID-MS/MS). The results of ^1^H–^13^C HSQC NMR experiments enabled us to assign ^1^H and ^13^C chemical shifts to seven distinct monosaccharides within the octasaccharide (Table S5); signals corresponding to the reducing-end glucosyl residue were not detected, presumably due to its anomeric heterogeneity. Cross-peaks produced in ^1^H-^1^H NMR experiments then confirmed the linkages between them (Fig. 4b). In particular, a cross-peak at (4.53, 3.62) ppm in NOESY experiments provided evidence of a β-xylosyl residue linked to the C2 hydroxyl of the second α-xylosyl residue of an XXXG structure. Furthermore, HPLC analysis of the monosaccharides released after total hydrolysis with trifluoroacetic acid (TFA) revealed that the octasaccharide is composed of a 1:1 ratio of xylose to glucose (Fig. 4c; exact ratio = (0.94 ± 0.04):1 Xyl:Glc as an average of three technical replicates). Finally, after derivatising the oligosaccharide with 2-aminobenzamide, CID-MS/MS confirmed that it possesses a tetrahexosyl backbone with a dipentosyl side chain attached to the second hexosyl unit and single pentosyl substitutions on the first and third (Fig. 4d). Therefore, the results of all experiments are consistent with the proposed XUXG (Xyl_4_Glc_4_) structure. Integrating data from the preceding experiments, the xyloglucan in *Cc*XBT1-(over)expressing plants is therefore likely made up predominantly of XUXG repeats in addition to a much smaller number of XUGG, XXXG, XUUG, and XUFG units.

### BoGH43A is a dual-function, promiscuous α-arabinofuranosidase/β-xylosidase

The presence of an apparent xyloglucan β-xylosyltransferase in *C. canephora*, from outside the Ericales order, suggests that β-xylosylated xyloglucan could be more widespread in the plant kingdom (and therefore the diet) than previously thought. Furthermore, the potential presence of other pentosyl xyloglucan decorations (such as the arabinopyranosylated D structure) has yet to be fully eliminated in higher plants. We were curious to see whether such structures can be metabolised by gut micro-organisms, as no xyloglucan β-xylosidases or α-arabinopyranosidases have yet been identified. Interestingly, the two *B. ovatus* xyloglucan α-arabinofuranosidases, *Bo*GH43A and *Bo*GH43B (encoded by neighbouring genes in the *B. ovatus* XyGUL; Figs. 5a,b), have markedly different levels of activity on α-arabinofuranosides, and it has been suggested that *Bo*GH43B may have evolved a different function to *Bo*GH43A (Larsbrink *et al*., 2014). Hence, we set out to analyse comprehensively the activities of the two enzymes.

**Figure 5.**
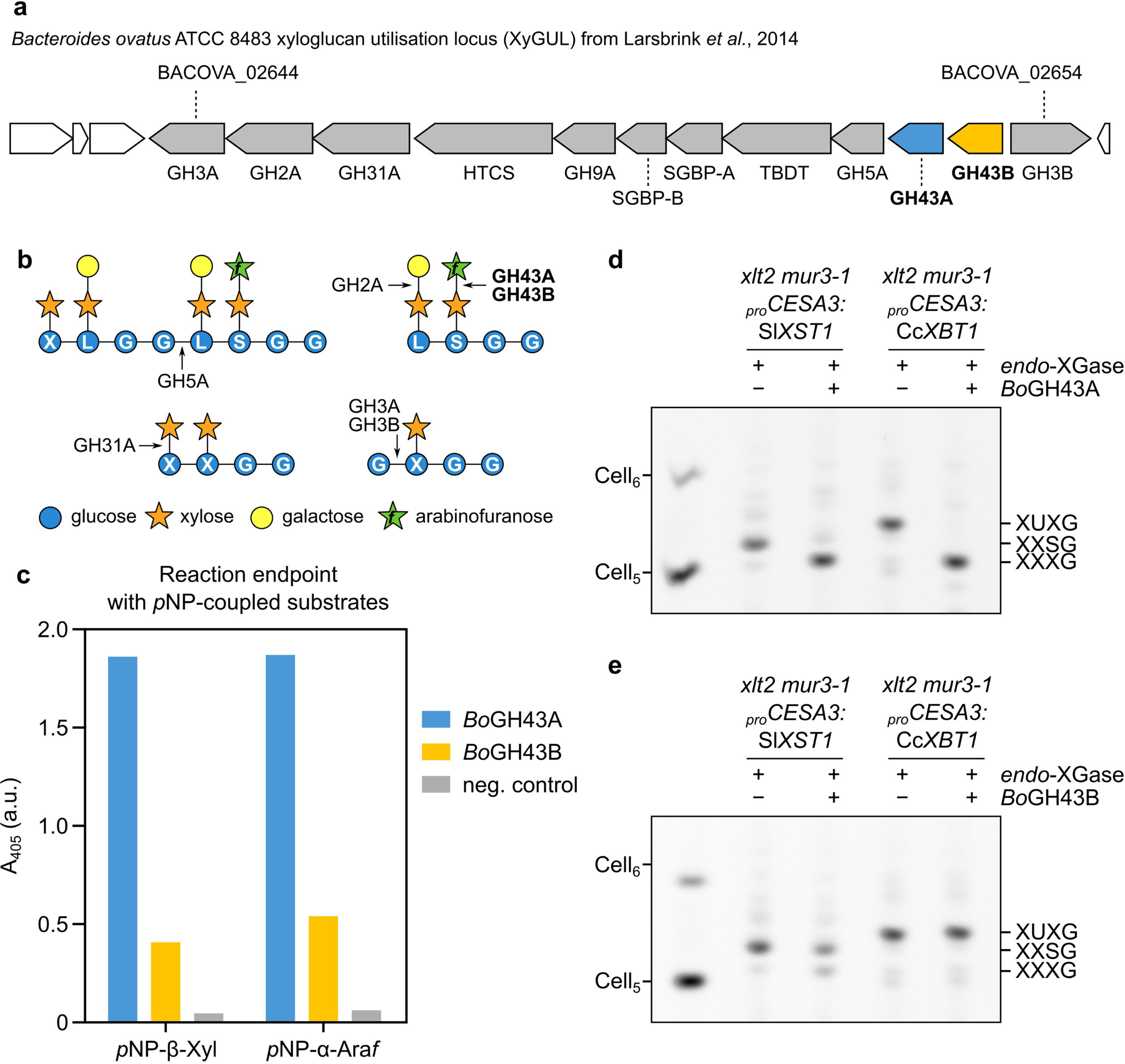
BoGH43A effectively hydrolyses both α-arabinosides and β-xylosides from synthetic and native substrates. **a** Schematic showing structure of xyloglucan utilisation locus (XyGUL) in the genome of *Bacteroides ovatus* ATCC 8483. *Bo*GH43A and *Bo*GH43B coding sequences are highlighted in blue and yellow, respectively. **b** Simplified schematic showing previously characterised XyGUL activities. **c** Activity test of *Bo*GH43A and *Bo*GH43B on *para*-nitrophenyl (*p*NP) glycosides. Absorbance values at A_405_ are endpoints measured after overnight incubation at pH 7.0. **d,e** Activity of *Bo*GH43A or *Bo*GH43B, respectively, on α-arabinosylated and β-xylosylated xyloglucan oligosaccharides produced from transgenic plants (PACE gels).

Kinetic data have already been collected for *Bo*GH43A on the synthetic substrates *para*-nitrophenyl α-arabinofuranoside (*p*NP-α-Ara*f*) and *p*NP-β-Xyl, whereas data on *Bo*GH43B have only been published for *p*NP-α-Ara*f* (Larsbrink *et al*., 2014). To obtain a more complete picture of their activities, we purified both enzymes (Fig. S7) and tested their activity on *p*NP-α-Ara*f*, *p*NP-β-Xyl, and *p*NP-α-Ara*p*. Both *Bo*GH43A and *Bo*GH43B were active on *p*NP-β-Xyl and *p*NP-α-Ara*f*, but not *p*NP-α-Ara*p,* with *Bo*GH43A appearing as a better catalyst for both substrates (Figs. 5c and S8), consistent with previous data (Larsbrink *et al*., 2014).

After determining the pH optimum for both enzymes at pH 7.5 in 50 mM HEPES buffer, we collected Michaelis-Menten kinetic data for both enzymes on *p*NP-α-Ara*f* and *p*NP-β-Xyl under these buffer conditions (Fig. S9). Similarly to previous results (Larsbrink *et al*., 2014), we found the catalytic efficiency (*k*_cat_/*K*_M_) of *Bo*GH43A for *p*NP-α-Ara*f* (20 M^-1^s^-1^) to be only twofold higher than that for *p*NP-β-Xyl (10 M^−1^s^−1^). This was in spite of much higher affinity for *p*NP-α-Ara*f* (K_M_ = 0.49 mM, *vs* 8.9 mM for *p*NP-β-Xyl), which was balanced out by a lower turnover number (*k*_cat_ = 0.01 s^−1^ *vs* 0.1 s^−1^ for *p*NP-β-Xyl). Although *Bo*GH43B exhibited catalytic parameters in the same order of magnitude for both substrates, they were almost two orders of magnitude lower than those of *Bo*GH43A (Table S6).

To ascertain whether *Bo*GH43A and *Bo*GH43B are active on *bona fide* β-xylosylated xyloglucan substrates, we incubated *endo*-XGase products from *Sl*XST1- and *Cc*XBT1-expressing plants with each enzyme in turn. Not only was *Bo*GH43A able to efficiently convert the α-arabinofuranosylated oligosaccharide XXSG to XXXG, but it was also able to convert β-xylosylated XUXG to XXXG—to at least the same, if not greater, completeness (Fig. 5d). By contrast, the same amount of *Bo*GH43B exhibited only a small amount of activity on the XXSG oligosaccharide and no detectable activity on XUXG (Fig. 5e). Hence, we were able to confirm that *Bo*GH43A, at least, can act both as a xyloglucan α-arabinofuranosidase and as a xyloglucan β-xylosidase.

### BoGH43A is active on naturally α-arabinofuranosylated and β-xylosylated xyloglucans, but fails to remove an unidentified pentosyl decoration from blueberry xyloglucan

To confirm that *Bo*GH43A can act on naturally β-xylosylated xyloglucan substrates, we also tested its ability to hydrolyse decorations from previously characterised arabinoxyloglucans and β-xylosylated xyloglucans from asterid cell walls. To simplify product characterisation, we focused on XXXG-type xyloglucans. Accordingly, we prepared AIR from the following four sources: (1) Arabidopsis leaf (lacking pentosyl decorations); (2) olive (*Olea europaea*, Lamiales, lamiids) leaf, since arabino/galacto-xyloglucan has been observed in olive fruits (Vierhuis *et al*., 2001a,b); (3) argan (*Argania spinosa*, Ericales) leaf, which exhibits a β-xylosylated fucogalactoxyloglucan (Ray *et al*., 2004); and (4) blueberry (*Vaccinium corymbosum*, Ericales) fruit skin, which was assumed to exhibit a similar β-xylosylated fucogalactoxyloglucan to that seen in bilberry (*V. myrtillus*) (Hilz *et al*., 2007). As before, alkali-extracted hemicellulose was cleaved into XyGOs by *endo*-XGase. The products were analysed by PACE (Figs. 6a and S10) and MALDI-TOF MS (Fig. S11), revealing that, as expected, the XyGOs mainly consisted of XXXG-based oligosaccharides with a wide range of decorations. We subjected these oligosaccharides to a series of combinatorial digestions with *Bb*AfcA α1,2-fucosidase, Fam35 β-galactosidase, and/or *Bo*GH43A. The identities of the products were confirmed with reference to their sensitivity to *Cj*Abf51 α1,2/3-arabinofuranosidase *vs Cg*GH3 β1,2-xylosidase (Fig. S12). Importantly, whereas *Bo*GH43A had no activity on Arabidopsis XyGOs (Fig. 6b), the enzyme was able to hydrolyse not only the α-arabinofuranosyl residues from olive-derived oligosaccharides XXSG, XLSG, and XSSG (Fig. 6c; increased enzyme concentration was required to complete the digestion), but also the β-xylosyl residues from argan and blueberry-derived XUXG and XUFG (Fig. 6d,e).

**Figure 6.**
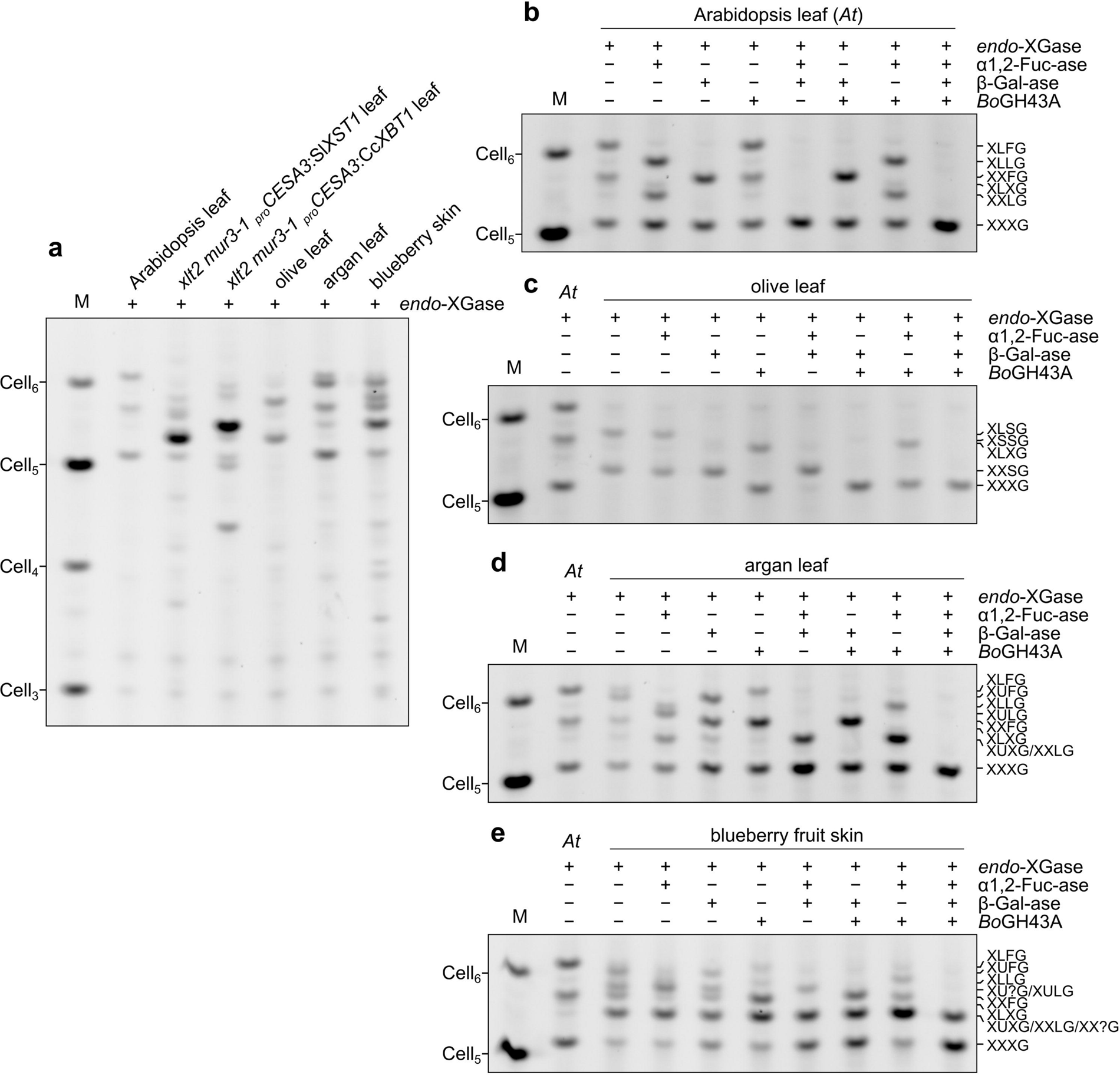
BoGH43A can assist in degrading both α-arabinosylated and β-xylosylated xyloglucans from natural sources. Various XXXG-type xyloglucans were extracted with alkali from AIR material. **a** PACE analysis of *endo*-XGase products without further digestion. See Fig. S10 for no-enzyme controls. **b–e** Combinatorial digest of *endo*-XGase products from **b** Arabidopsis leaf xyloglucan, **c** olive (*Olea europaea*) leaf xyloglucan, **d** argan (*Argania spinosa* / *Sideroxylon spinosum*) leaf xyloglucan, or **e** blueberry (*Vaccinium corymbosum*) fruit skin xyloglucan with *Bb*AfcA α1,2-fucosidase (α1,2-Fuc-ase), Fam35 β-galactosidase (β-Gal-ase), and/or *Bo*GH43A (PACE gels). Assignments in all panels were assisted with MALDI-TOF mass spectrometry of the *endo*-XGase products (Fig. S11).

However, interestingly, one of the blueberry XyGOs—which co-migrated with XUXG and XXLG in PACE analysis—appeared to be completely resistant to all three enzymes (Fig. 6e). We enriched this oligosaccharide by size exclusion but were unable to purify it sufficiently for NMR characterisation. Nevertheless, CID MS–MS experiments indicated that it its structure is likely XXXG with a pentosyl decoration on the third xylose (Fig. S13). A β1,2-linked xylosyl decoration on the third xylose has been previously reported in bilberry (Hilz *et al*., 2007); hence, we hypothesised that the recalcitrant blueberry XyGO might be XXUG, and that *Bo*GH43A might not tolerate the structure of this oligosaccharide. However, the recalcitrant blueberry XyGO was also resistant to *Cg*GH3 β1,2-xylosidase (as well as to *Cj*Abf51 α1,2/3-arabinofuranosidase; Fig. S12c). Furthermore, removal of the non-reducing terminal isoprimeverosyl unit by sequential digestion with α-xylosidase and β-glucosidase did not alter its insensitivity to *Bo*GH43A (Fig. S14). Although more work will be required to identify this oligosaccharide, our results suggest that blueberry xyloglucan could be decorated with a hitherto unknown xyloglucan side chain—perhaps even the speculative α-xylosylated xylose disaccharide that was previously reported in tobacco and aubergine (Sims *et al*., 1996; Kato *et al*., 2010). These results highlight the importance of using comprehensive glycosidase- and/or NMR-based experiments for characterising xyloglucan structure, as opposed to assignments based on mass spectrometry alone.

### β-Xylosylated xyloglucan may be restricted to specific, undetermined tissues in coffee and other asterid species

Since β-xylosylated xyloglucan has so far been detected in only two species from the Ericales order, we were also prompted to investigate the true prevalence of β-xylosylated xyloglucan in asterids. Surprisingly, despite the presence of *Cc*XBT1 in the *C. canephora* genome, our methods were unable to detect β-xylosylation of xyloglucan extracted from leaves of the lamiids *C. arabica* (Arabica coffee; Figs. S15, S16), *Catharanthus roseus* (Gentianales; Fig. S17), or *Borago officinalis* (borage, Boraginales; data not shown). However, we also investigated the structure of xyloglucan from the exocarp of kiwi (*Actinidia chinense*), a member of the Ericales. *Endo*-XGase products from kiwi endocarp exhibited a similar size distribution to those from tomato fruit, which is known to exhibit an XXGG-type arabinoxyloglucan (Fig. S18a). Unlike tomato XyGOs, however, kiwi XyGOs were sensitive to *Cg*GH3 β1,2-xylosidase and *Bo*GH43A, but not *Cj*Abf51 α1,2/3-arabinofuranosidase (Figs. 7a,b and S18b). With aid from MALDI-TOF MS analysis of untreated and *Bo*GH43A-treated *endo*-XGase products (Fig. 7c), we assigned the main two XyGOs as XUGG and XUG (the latter likely arising from alternative cleavage of the deacetylated backbone by *endo*-XGase). Hence, kiwi fruit skin xyloglucan is likely composed mainly of repeating units of XUGG, making it unique among currently characterised xyloglucan structures.

**Figure 7.**
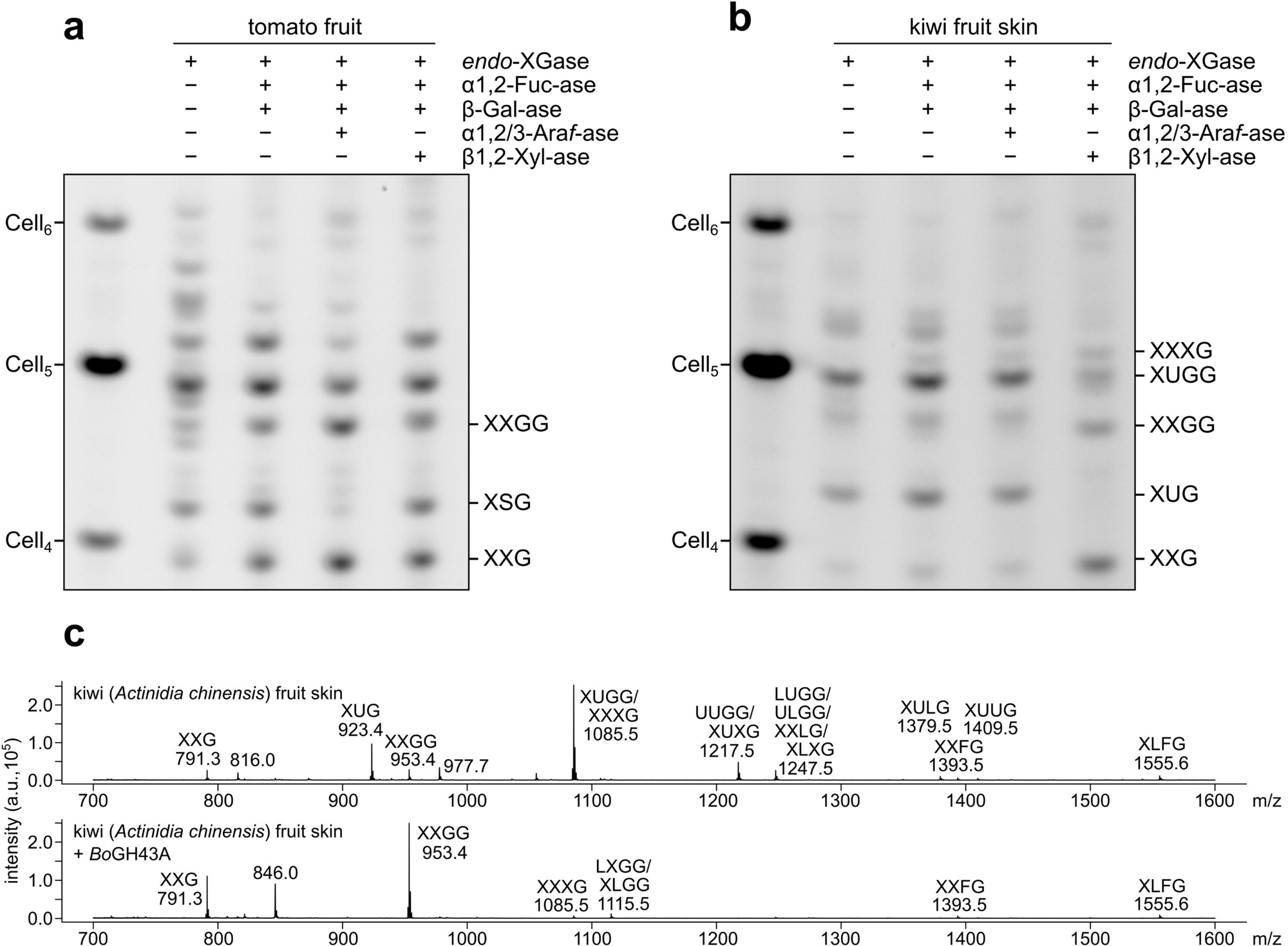
Xyloglucan from kiwi fruit skin appears to exhibit a xyloglucan made up predominantly of XUGG units. Alkali-extracted hemicellulose from green tomato fruit or kiwi fruit skin was digested with *Aa*XEG *endo*-XGase followed by ethanol precipitation. **a** Combinatorial digest of *endo*-XGase products from tomato with *Bb*AfcA α1,2-fucosidase (α1,2-Fuc-ase), Fam35 β-galactosidase (β-Gal-ase), *Cj*Abf51 α1,2/3-arabinofuranosidase (α1,2/3-Ara*f*-ase), and/or *Cg*GH3 β1,2-xylosidase (PACE gel). **b** Combinatorial digest of *endo*-XGase products from kiwi (PACE gel). See also the similar results in Fig. S18 from combinatorial digestions involving *Bo*GH43A. The assignment of the band labelled XXG in **a** and **b** is based on co-migration with the XXG product seen in sequential digest experiments on XXXG-type xyloglucan (e.g. Fig. S14). Assignments of XSG and XXGG are tentative but deduced from the enzyme activities and expected band shifts (to XXG or from XUGG, respectively). Assignments of kiwi XyGOs are consistent with mass spectrometry results. **c** MALDI-TOF mass spectrometry analysis of *endo*-XGase products from kiwi treated with or without *Bo*GH43A.

## Discussion

It was initially speculated that the β-xylosyl residues of argan leaf xyloglucan might be the result of a galactosyltransferase side-activity (Ray *et al*., 2004). However, in this work, we have identified a dedicated xyloglucan β-xylosyltransferase (*Cc*XBT1), which, when expressed in Arabidopsis, is specific to the second xylosyl residue in the XXXG repeat. Interestingly, the presence of XUGG and XUG *endo*-XGase products from Arabidopsis plants over-expressing *Cc*XBT1 suggests that the high level of *Cc*XBT1 activity might somehow alter the α-xylosylation pattern laid down by XXTs in the Golgi. We speculate that early modification of the second α-xylosyl residue by the over-expressed *Cc*XBT1 enzyme might inhibit the addition of the third.

Expression of *Cc*XBT1 in Arabidopsis galactosyltransferase mutants *mur3-3* and *xtl2 mur3-1* rescued growth, revealing that, like arabinoxyloglucan (Schultink *et al*., 2013), β-xylosylated xyloglucan can functionally replace fucogalactoxyloglucan in Arabidopsis cell biology. Furthermore, the fact that (over-)expression of either *At*XLT2 (Kong *et al*., 2015) or *Cc*XBT1 (which are both strongly specific for the second α-xylosyl residue) can rescue growth of *mur3-3* and *xtl2 mur3-1* demonstrates unequivocally that decoration of the third α-xylosyl residue is not necessary for normal growth in Arabidopsis. Therefore, it seems that it is undecorated xyloglucan in general that is somehow dysfunctional or toxic. This is consistent with the previous proposal that undecorated xyloglucan aggregates in the Golgi, preventing its secretion (Tamura *et al*., 2005; Kong *et al*., 2015).

In addition to *Cc*XBT1, we also identified a putative xyloglucan arabinofuranosyltransferase from cranberry (*Vaccinium macrocarpon*, Ericales): *Vm*XST1. The activity of this enzyme is consistent with previous reports that arabinofuranosylated XyGOs can be directly extracted from cranberry fruits (Hotchkiss *et al*., 2015; Sun *et al*., 2015; Auker *et al*., 2019). When expressed in Arabidopsis, *Vm*XST1 exhibited only a very small amount of activity; the reason for this is not clear, but it is possible that, *in planta*, it is redundant to other, more active paralogues. Nevertheless, the expression of *Vm*XST1 in Arabidopsis galactosyltransferase mutants afforded substantial (albeit incomplete) complementation, suggesting that the level of xyloglucan decoration needs only to reach a low threshold to permit normal growth. Importantly, this result suggests that the fundamental cell biological properties of xyloglucan are unlikely to be the driving force for xyloglucan side chain diversity.

*Cc*XBT1 and *Vm*XST1 are members of GT47-A, as was originally proposed for the Ericales β-xylosyltransferase by Schultink et al. (2014). More specifically, these enzymes are members of GT47-A_III_ (alongside *At*XLT2, *Os*XLT2, *Sl*XST1, and *Sl*XST2). Here, we have comprehensively mapped the expansion of this clade in asterid genomes, establishing the presence of as many as four paralogues (*a*, *b*, *c*1, and *c*2) in many lamiid genomes. Our analysis grouped *Cc*XBT1 in GT47-A_III_*c*2 together with orthologues from many other lamiid species, including olive and *C. roseus*. However, we were unsuccessful in detecting β-xylosylated xyloglucan in any lamiid; hence, it is unclear whether the β-xylosyltransferase activity is unique to *Cc*XBT1 or whether β-xylosylated xyloglucan is restricted to a particular organ or tissue in these species. Precedence for the latter lies in the fact that XXXG-type fucogalactoxyloglucan has only been found in the roots of grasses and the pollen tubes of Solanaceae-family plants (Lampugnani *et al*., 2013; Dardelle *et al*., 2015; Liu *et al*., 2015); similarly, Arabidopsis galacturonoxyloglucan appears limited to root hairs (Peña *et al*., 2012). A third explanation could be that β-xylosylated xyloglucan is for some reason difficult to extract from the cell walls of these species.

Our data suggest that changes to certain residues surrounding the donor sugar binding site may explain the apparent neofunctionalisation seen in GT47-A_III_*c*. Despite this, we were unable to identify a compelling pattern relating the observed activities to the corresponding protein sequences. Although studying the evolution of these sites could still accelerate the discovery of new and unexpected activities, it is possible that GT47-A nucleotide sugar specificity is too subtle to be predicted from amino acid identities alone; characterisation of further GT47-A_III_ activities and point mutation experiments will be required to confirm this. In particular, we suggest that the distinction between UDP-β-L-Ara*f* and UDP-α-D-Xyl may be especially subtle. This could explain the apparent rapid fluctuations between arabinofuranosyltransferase and xylosyltransferase activity as illustrated in Fig. 2a. Supporting this idea, the same phenomenon has been observed in the unrelated GT61 family (Cenci *et al*., 2018; Zhong *et al*., 2022). Interestingly, however, and in contrast to glycosyl hydrolases, there is not yet any evidence for Ara*f*/Xyl promiscuity in these families.

Many species in the Ericales order produce commonly consumed fruits, nuts, and leaves; hence, β-xylosylated xyloglucan is likely a substantial component in human diets. Here, we have revealed the probable pathway for its disassembly in *B. ovatus* and similar species, which are thought to convert xyloglucan into beneficial short-chain fatty acids (Cantu-Jungles *et al*., 2019; Liu *et al*., 2020). Although we initially hypothesised that *Bo*GH43B might have a novel activity, we found that *Bo*GH43A is probably the major enzyme responsible for processing both arabinoxyloglucan and β-xylosylated xyloglucan. Similar promiscuity is also found in related GH43-family xylan α-arabinosidases/β-xylosidases (Rogowski *et al*., 2015), suggesting that α-L-arabinosides and β-D-xylosides could also have similar chemical properties. However, although *Bo*GH43B exhibited promiscuity for the two *p*NP-labelled sugars (Figs. 5c, S8, and S9), it did not exhibit activity on the β-xylosylated XUXG oligosaccharide (Figs. 5d,e). This discrepancy might stem from differences in substrate concentration or fundamental kinetic parameters between the model and native substrates. Nevertheless, it cannot yet be eliminated that some xyloglucan GH43 enzymes are specific for one of the two sugars; for instance, the putative xyloglucan α-arabinofuranosidase *Xac*Abf43A from *X. citri* pv. *citri* exhibits tenfold lower activity on *p*NP-Xyl than on *p*NP-Ara*f* (Vieira *et al*., 2021). This could provide an incentive for plants to develop more diverse xyloglucan structures as a means of pathogen resistance. In turn, such adaptations likely incentivise promiscuity in microbial cell wall-degrading enzymes.

To conclude, our results not only expand the existing knowledge of xyloglucan diversity, but also, by identifying an enzyme capable of adding β-xylosyl decorations, make a significant step towards understanding and predicting it from sequence level. In terms of methodology, the glycosidase assays and PACE experiments that we present here also represent the most comprehensive system for analysing xyoglucan structure to date. Combined with knowledge of microbial metabolism in the gut, these results will ultimately allow us to select the best dietary fibres for a balanced gut microbiome and thereby empower us to improve our digestive health.

## Acknowledgements

This work was supported by a grant to P.D. from OpenPlant (BB/L014130/1, P.D.) and a grant to F.H. from the ERC (695669). L.F.L.W. was supported by the University of Cambridge, Jesus College, Cambridge, and the Department of Plant Sciences, Cambridge. We thank Drs Xiaolan Yu, Henry Temple, and Yoshihisa Yoshimi (University of Cambridge) for technical assistance, as well as Ms Mar Millan and Mr Alex Summers (Cambridge University Botanic Garden) for assistance in obtaining plant material, and Profs. Harry Brumer (University of British Columbia) and Gideon Davies (University of York) for providing expression constructs for *Bo*GH43A and *Bo*GH43B. We also thank Mr Konan Ishida (University of Cambridge) for critical reading of the manuscript.

## Author contribution

LFLW, SN, KS, and PD designed experiments. LFLW conducted bioinformatic analyses and molecular phylogenies, performed molecular biology, and generated all plant genotypes. SN performed protein purification and colorimetric assays. LFLW and SN performed glycosidase digestions and PACE assays. LFLW, LY, and SN conducted oligosaccharide gel filtration. LY and KS carried out NMR analysis. TT and LFLW performed mass spectrometry. PD and FH supervised LFLW and SN, respectively. LFLW wrote the manuscript, to which SN and all other authors also contributed.

