## Supporting Information for "The biosynthesis, degradation, and function of cell wall β-xylosylated xyloglucan mirrors that of arabinoxyloglucan"

The following Supporting Information is available for this article:

**Fig. S1** Unrooted maximum-likelihood phylogeny of GT47-A sequences from a range of asterid and non-asterid species, showing recent expansion of the GT47-AIII clade in asterids.

**Fig. S2** GT47-$A_{III}$ subtree from Fig. 2 and Fig. S1, fully labelled.

**Fig. S3** Reported confidence metrics for the AlphaFold model of *At*XLT2.

**Fig. S4** Individual GT47-A $T_1$ transformants in *mur3-3* background.

**Fig. S5** Individual GT47-A $T_1$ transformants in *xlt2 mur3-1* background.

**Fig. S6** No-enzyme controls for *endo*-XGase digestions of transgenic Arabidopsis material.

**Fig. S7** SDS-PAGE gels of purified *Bo*GH43A and *Bo*GH43B.

**Fig. S8** Initial timecourses of *Bo*GH43A and *Bo*GH43B activity on *para*-nitrophenyl (*p*NP) glycosides *p*NP-α-Ara*f*, *p*NP-β-Xyl, and *p*NP-α-Ara*p* in order to screen for potential previously unreported activities.

**Fig. S9** pH optima and Michaelis-Menten curves for *Bo*GH43A and *Bo*GH43B activity on *para*-nitrophenyl (*p*NP) glycosides.

**Fig. S10** No-enzyme controls for *endo*-XGase digestions of plants exhibiting XXXG-type xyloglucan.

**Fig. S11** MALDI-TOF mass spectrometry analysis of *endo*-xyloglucanase products from olive leaf, argan leaf, and blueberry fruit skin.

**Fig. S12** Sensitivity of olive, argan, and blueberry *endo*-XGase products to α1,2/3-arabinofuranosidase and β1,2-xylosidase.

**Fig. S13** Spectrum from tandem mass spectrometry (MS–MS) with collision-induced dissociation (CID) of uncharacterised blueberry xyloglucan oligosaccharide.

**Fig. S14** Sequential digestion of an unidentified *endo*-XGase product from blueberry skin xyloglucan.

**Fig. S15** Partial characterisation of *endo*-XGase products from *Coffea arabica* leaf xyloglucan.

**Fig. S16** No-enzyme controls for *endo*-XGase digestions of plants exhibiting XXGG-/XXG$_n$-type xyloglucan.

**Fig. S17** Sensitivity of *endo*-XGase products from *Catharanthus roseus* leaf xyloglucan to various *exo*-glycosidases.

**Fig. S18** Characterisation of *endo*-XGase products from kiwi fruit skin xyloglucan.

**Table S1** Genomic data sources.

**Table S2** De novo-synthesised coding sequences for Golden Gate assembly.

**Table S3** PCR primers used to amplify DNA parts for Golden Gate assembly.

**Table S4** *Exo*-glycosidases used in this work.

**Table S5** $^{1}$H and $^{13}$C NMR assignments for XUXG oligosaccharide.

**Table S6** Kinetic parameters for *Bo*GH43A/B activity on *para*-nitrophenyl glycosides.

**Methods S1** Glycosyl hydrolase expression and purification.

**Methods S2** Oligosaccharide purification.

**Fig. S1 Unrooted maximum-likelihood phylogeny of GT47-A sequences from a range of asterid and non-asterid species, showing recent expansion of the GT47-$A_{III}$ clade in asterids.** See Fig. S2 or Table S1 for a full list of species. Known donor substrate specificities are annotated with sugar symbols (as designated in the key). Sequences were truncated to the GT47 domain prior to phylogenetic inference using IQ-TREE. Branch lengths indicate average number of subsititutions per site (refer to scale bar). Support values at important splits represent percentage replication within 1,000 ultra-fast bootstrap pseudo-replicates. GT47-A subclades are annotated with roman numerals (I–VII) as in Yu *et al*. (2022). GT47-A subclade III (GT47-$A_{III}$) is highlighed in grey; lower level subclades within GT47-$A_{III}$ (*a*, *b*, *c* etc.) are also annotated. Since *Sl*MUR3 is annotated as two separate loci in Phytozome (Solyc09g064470.3.1 and Solyc09g064480.1.1), its sequence was automatically was filtered out by the length threshold imposed during quality control. However, our previous phylogeny (Yu *et al*., 2022; DOI: 10.1093/plcell/koac238) grouped it in GT47-$A_{VI}$.

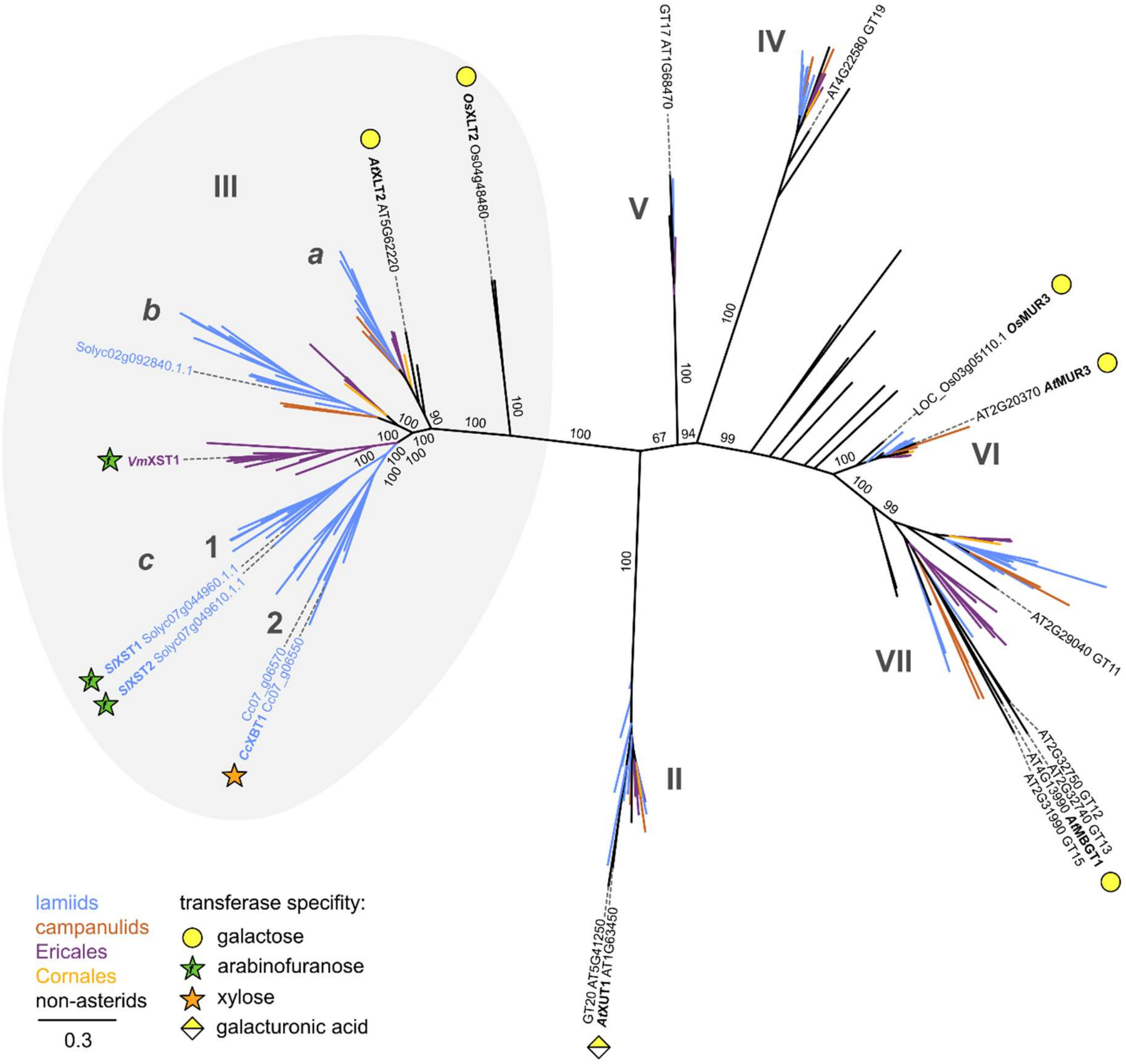

III
a
b
c
1
2
IV
V
VI
VII
II
OsXLT2 Os04g48480
AtXLT2 AT5G62220
Solyc02g092840.1.1
VmXST1
SlXST1 Solyc07g044960.1.1
SlXST2 Solyc07g049610.1.1
Cc07_g06570
CcXBT1 Cc07_g06550
GT17 AT1G68470
AT4G22580 GT19
LOC_Os03g05110.1 OsMUR3
AT2G20370 AtMUR3
AT2G29040 GT11
AT2G32750 GT12
AT2G32740 GT13
AT4G13990 AtMBGT1
AT2G31990 GT15
GT20 AT5G41250
AtXUT1 AT1G63450
lamiids
campanulids
Ericales
Cornales
non-asterids
0.3
transferase specifity:
galactose
arabinofuranose
xylose
galacturonic acid

**Fig. S2 GT47-$A_{III}$ subtree from Fig. 2 and Fig. S1, fully labelled.** Horizontal branch lengths indicate average number of subsititutions per site (refer to scale bar). Support values at each split represent percentage replication within 1,000 ultra-fast bootstrap pseudo-replicates. See Table S1 for full information on the source of the sequences.

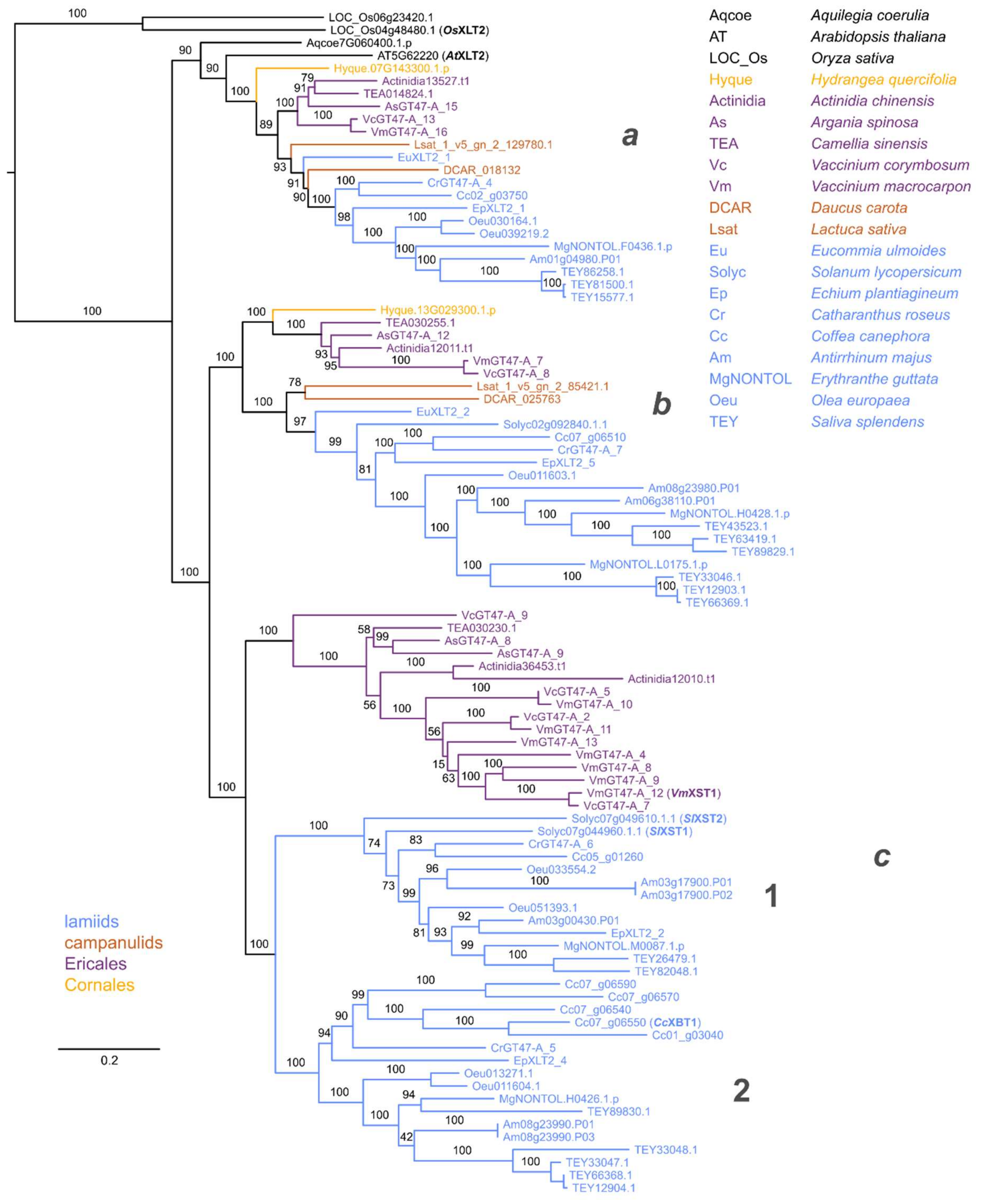

**Fig. S3 Reported confidence metrics for the AlphaFold model of *At*XLT2.** Figures were generated at the AlphaFold Protein Structure database (https://alphafold.ebi.ac.uk). **a** Per-residue confidence scores representing the predicted local distance difference test (pLDDT) for full-length *At*XLT2, including transmembrane helix, stem domain, and catalytic domain. Residues with pLDDT < 50 are predicted to be disordered. **b** Predicted aligned error plot.

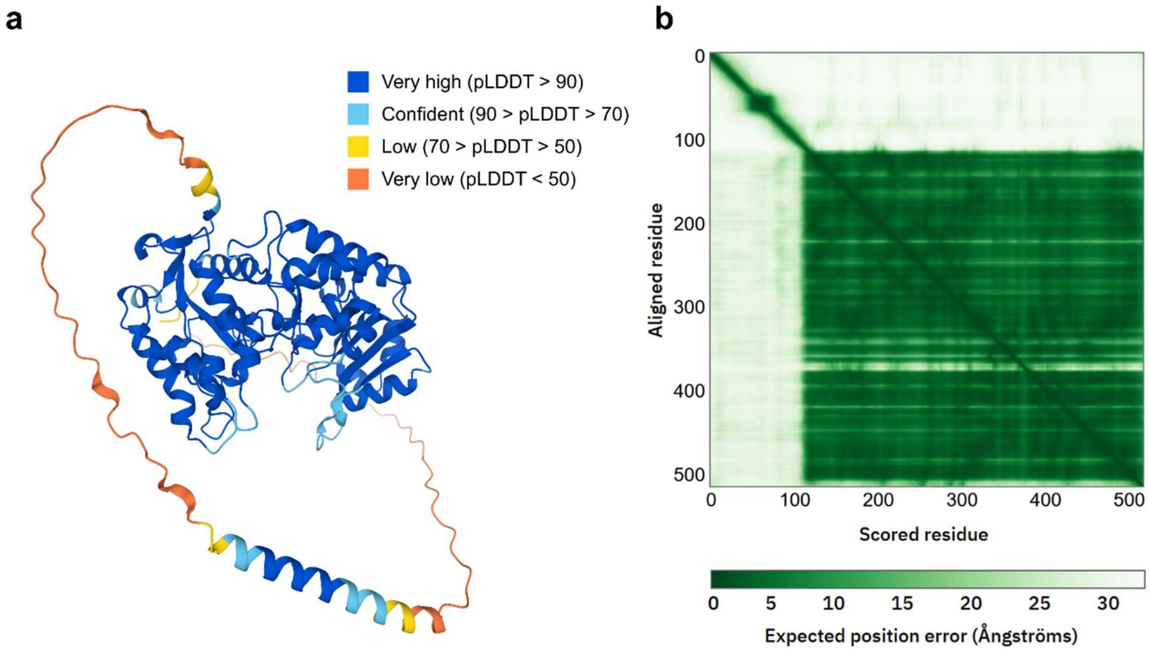

**Fig. S4 Individual GT47-A $T_1$ transformants in *mur3-3* background.** Plants were photographed after six weeks of growth. For the transgenic plants, each plant is a separate transgenic line (lines are labelled #1, #2, etc.). Transgenes were expressed under the promoter of xyloglucan α-xylosyltransferase XXT2. Expression of *Cc*XBT1 resulted in full rescue of *mur3-3*'s stunted growth, whereas expression of *Vm*XST1 afforded only partial complementation. Transformation with Cc07_g06570 had no effect apparent on the phenotype of *mur3-3*.

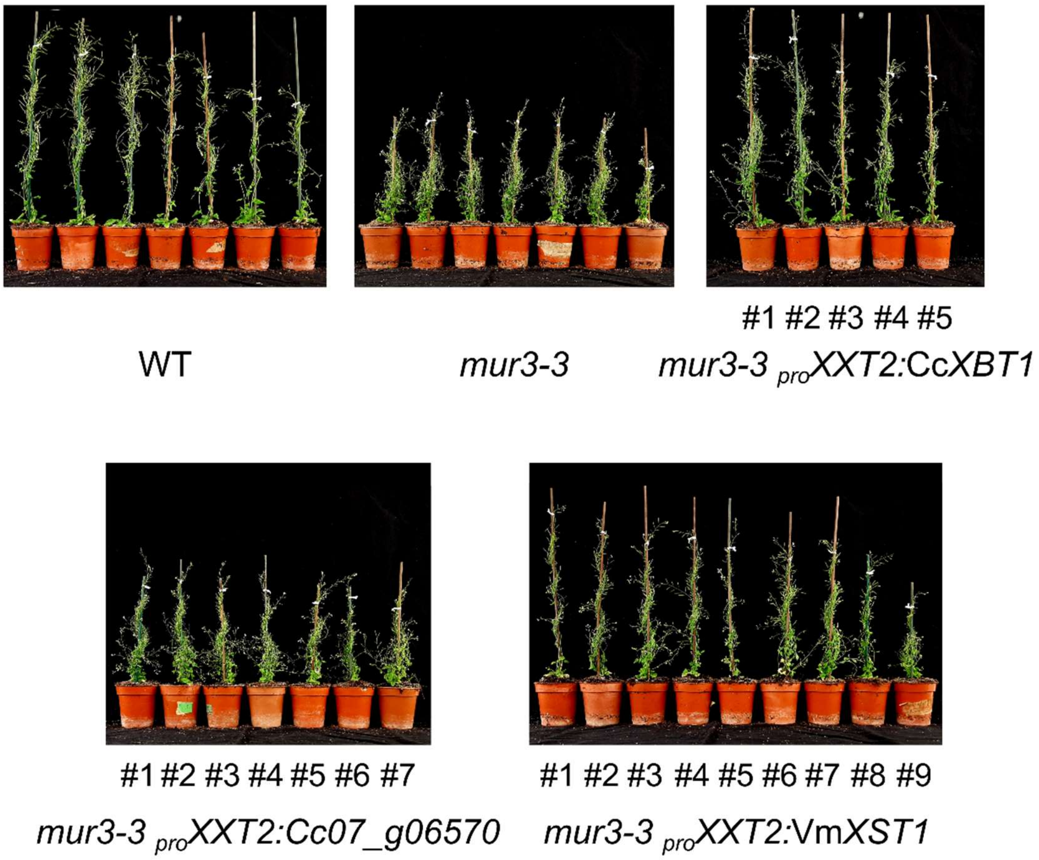

**Fig. S5 Individual GT47-A $T_1$ transformants in *xlt2 mur3-1* background. a** Plant growth phenotypes. Plants were photographed after six weeks of growth. For the transgenic plants, each plant is a separate transgenic line. **b** PACE analysis of *endo*-xyloglucanase (*endo*-XGase) products from transgenic cell wall material. Hemicellulose was extracted by alkali treatment of leaf alcohol insoluble residue (AIR) and digested with *Aa*XEG *endo*-XGase. Products were subsequently derivatised with a fluorophore and separated by electrophoresis. Individual transgenic lines are numbered in accordance with the photographs in panel **a**.

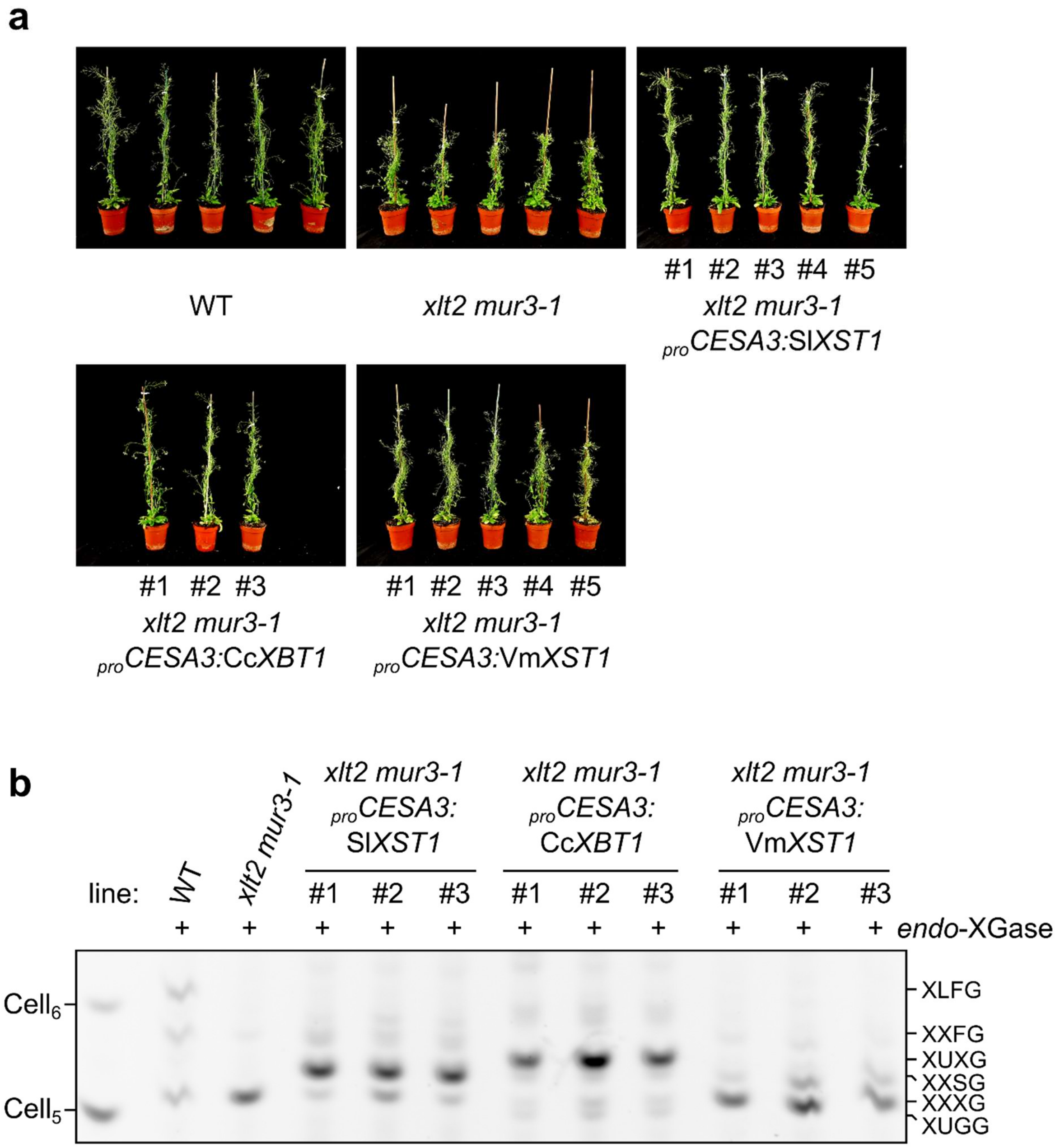

**Fig. S6 No-enzyme controls for *endo*-XGase digestions of transgenic Arabidopsis material.** Alkali-extracted hemicellulose from transgenic plants was treated with or without *Aa*XEG *endo*-XGase to confirm that the bands displayed in Fig. 3 result from specific hydrolysis by *endo*-XGase, as opposed to background contamination. Products were subsequently derivatised with a fluorophore and separated by electrophoresis.

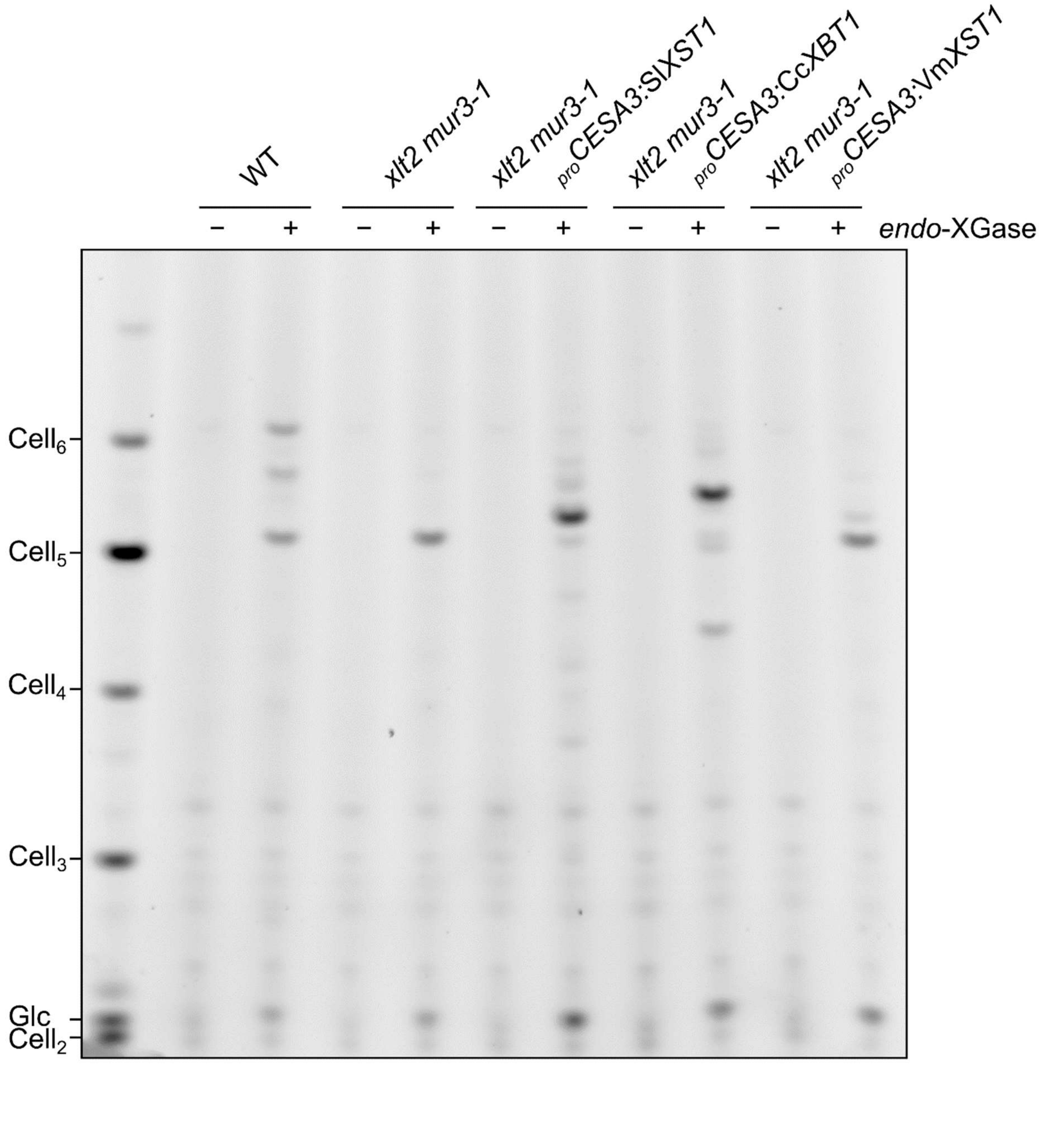

**Fig. S7 SDS-PAGE gels of purified *Bo*GH43A and *Bo*GH43B.** Purified proteins migrated consistently with their theoretical molecular weight (*Bo*GH43A: 57.6 kDa; *Bo*GH43B: 57.2 kDa).

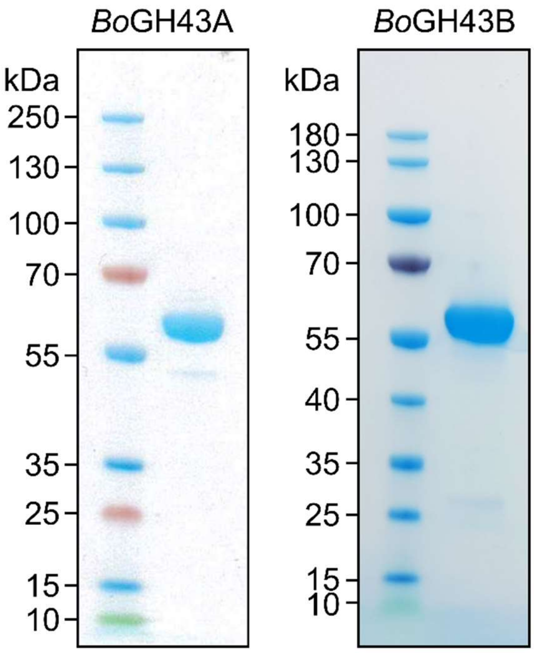

**Fig. S8 Initial timecourses of *Bo*GH43A and *Bo*GH43B activity on *para*-nitrophenyl (*p*NP) glycosides *p*NP-α-Ara*f*, *p*NP-β-Xyl, and *p*NP-α-Ara*p* in order to screen for potential previously unreported activities.** *p*NP glycosides were incubated at 1 mM concentration with 2 µM enzyme in 50 mM buffer, pH 7.0, T = 25°C. Reaction progress was monitored by observing $A_{405}$ at 1 min intervals. The lack of hydrolysis of *p*NP-α-Ara*p* even at 1 mM substrate concentration indicates a lack of activity and therefore the kinetics were not investigated further.

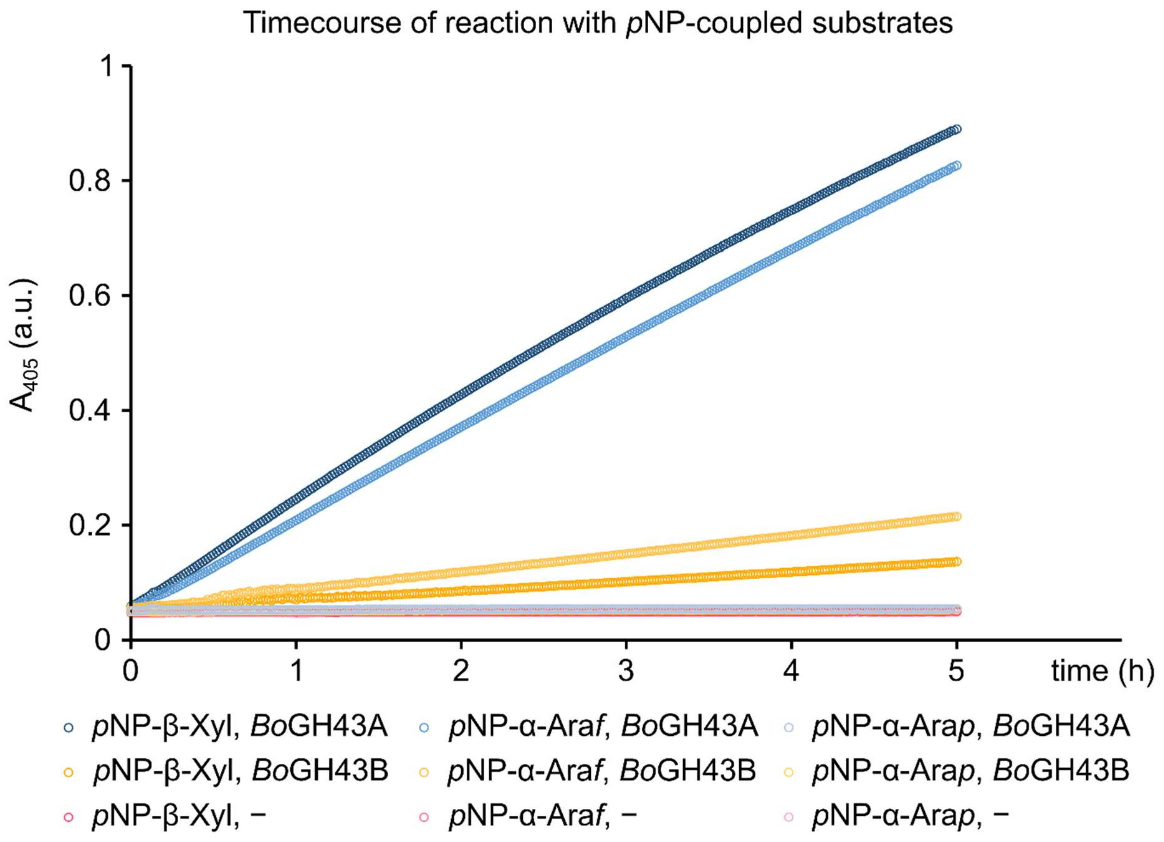

**Fig. S9 pH optima and Michaelis-Menten curves for *Bo*GH43A and *Bo*GH43B activity on *para*-nitrophenyl (*p*NP) glycosides. a,b** Determination of pH optima. Initial reaction velocities of **a** *Bo*GH43A on *p*NP-β-Xyl and **b** *Bo*GH43B on *p*NP-α-Ara*f*, determined by monitoring $A_{405}$ with 1 mM substrate and 2 µM enzyme in 50 mM buffer. Note that neither enzyme has any activity in Tris-HCl buffer. Tris has been observed to bind strongly to the active site of *Bo*GH43A in crystal structures (Hemsworth *et al.*, 2016). **c–f** Kinetic curves for *Bo*GH43A and *Bo*GH43B activity on *p*NP-α-Ara*f* (**c,d**) and *p*NP-β-Xyl (**e,f**). All kinetic measurements were carried out with 1 µM enzyme in 50 mM HEPES pH 7.5 at 20 °C. Reactions were monitored with $A_{405}$ in a plate reader. Kinetic data are summarized in Table S6.

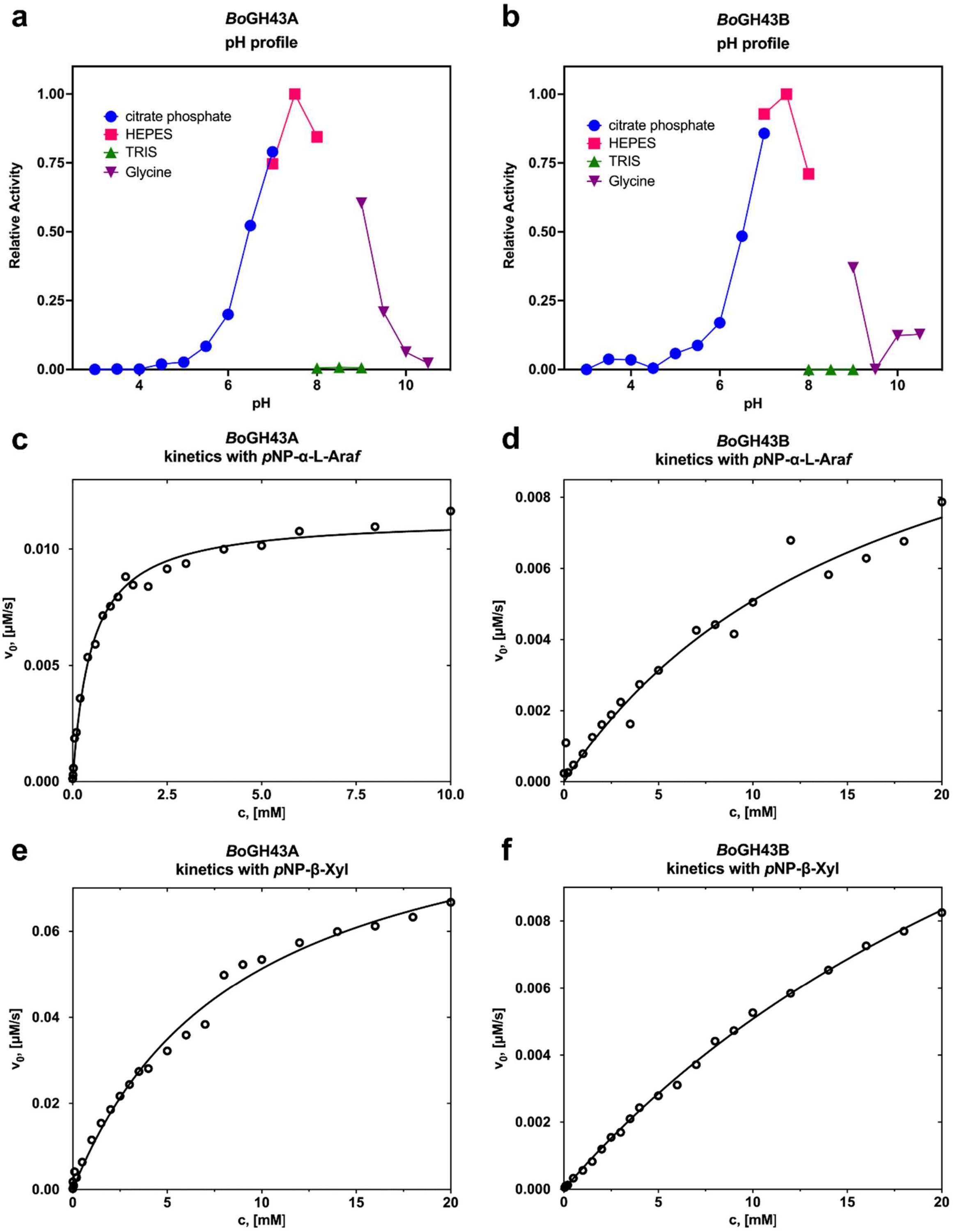

a
BoGH43A
pH profile
Relative Activity
citrate phosphate
HEPES
TRIS
Glycine
pH
b
BoGH43B
pH profile
c
BoGH43A
kinetics with pNP-α-L-Araf
v0, [µM/s]
c, [mM]
d
BoGH43B
kinetics with pNP-α-L-Araf
e
BoGH43A
kinetics with pNP-β-Xyl
f
BoGH43B
kinetics with pNP-β-Xyl

**Fig. S10 No-enzyme controls for *endo*-XGase digestions of plants exhibiting XXXG-type xyloglucan.** Alkali-extracted hemicellulose from transgenic plants was treated with or without *Aa*XEG endo-XGase to confirm that the bands displayed in Fig. 6 result from specific hydrolysis by *endo*-XGase, as opposed to background contamination. Products were subsequently derivatised with a fluorophore and separated by electrophoresis.

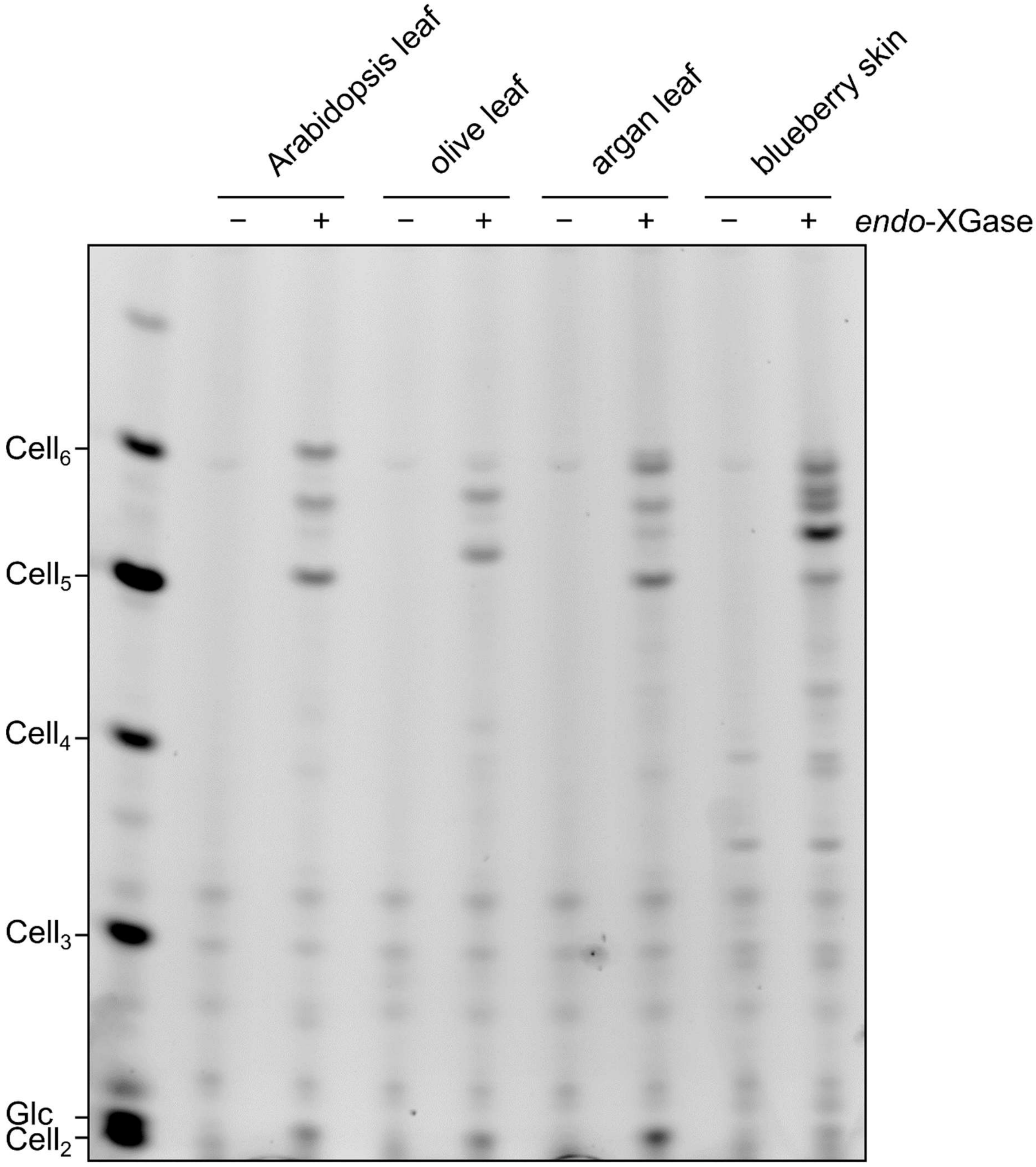

**Fig. S11 MALDI-TOF mass spectrometry analysis of *endo*-xyloglucanase products from olive leaf, argan leaf, and blueberry fruit skin.** Alkali-extracted hemicellulose was treated with *Aa*XEG *endo*-XGase. H = unknown hexose; P = unknown pentose.

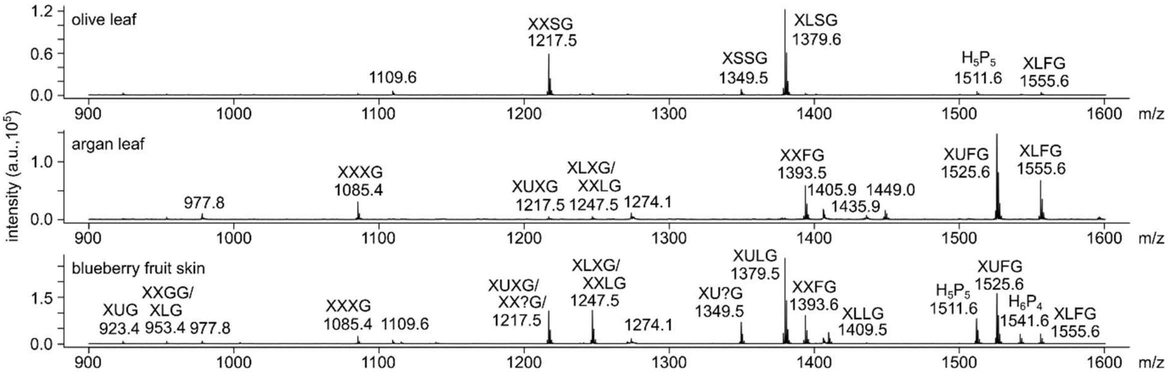

**Fig. S12 Sensitivity of olive, argan, and blueberry *endo*-XGase products to α1,2/3-arabinofuranosidase and β1,2-xylosidase.** Alkali-extracted hemicellulose from green tomato fruit or kiwi fruit skin was digested with *Aa*XEG *endo*-XGase before being subjected to ethanol precipitation. Oligosaccharides were then treated to combinatorial digestion with *Bb*AfcA α1,2-fucosidase (α1,2-Fuc-ase), Fam35 β-galactosidase (β-Gal-ase), *Cj*Abf51 α1,2/3-arabinofuranosidase (α1,2/3-Ara*f*-ase), and/or *Cg*GH3 β1,2-xylosidase. Products were subsequently derivatised with a fluorophore and separated by electrophoresis. **a** Digestion of *endo*-XGase from olive leaf. **b** Digestion of *endo*-XGase from argan leaf. **c** Digestion of *endo*-XGase from blueberry fruit skin.

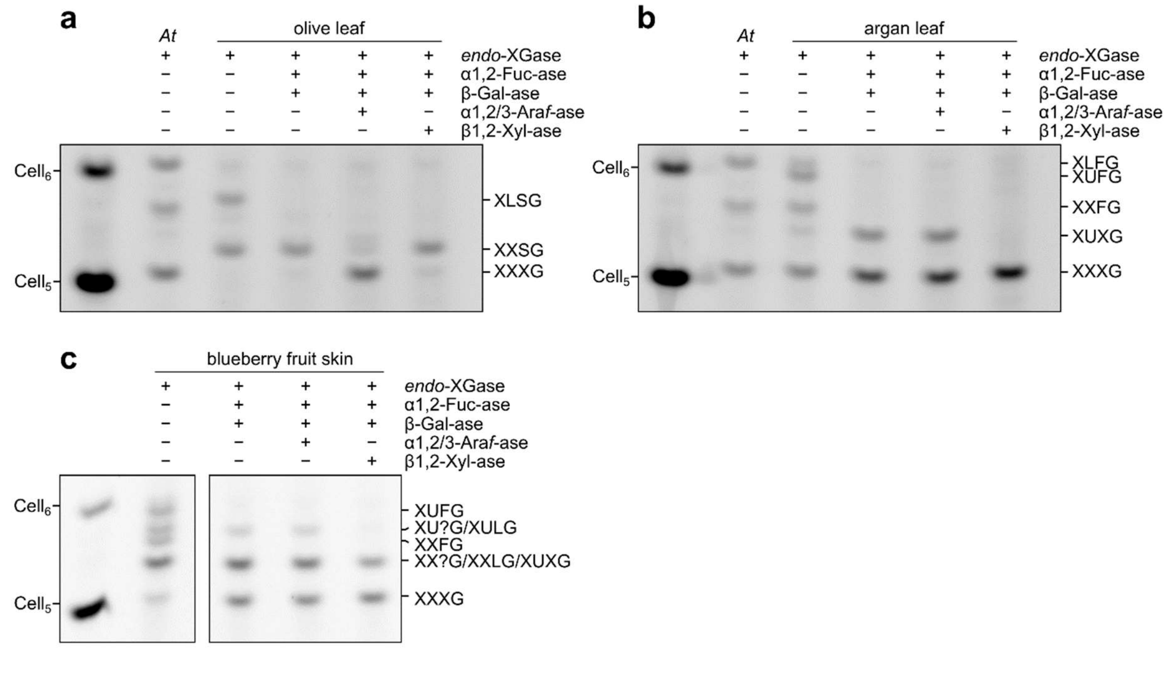

**Fig. S13 Spectrum from tandem mass spectrometry (MS–MS) with collision-induced dissociation (CID) of uncharacterised blueberry xyloglucan oligosaccharide.** The unidentified octasaccharide was semi-purified from alkali-treated blueberry skin AIR by treatment of *Aa*XEG *endo*-XGase products with *Bb*AfcA α1,2-fucosidase, Fam35 β-galactosidase, and *Bo*GH43A followed by size exclusion chromatography and reducing-end derivatisation with 2-aminobenzamide.

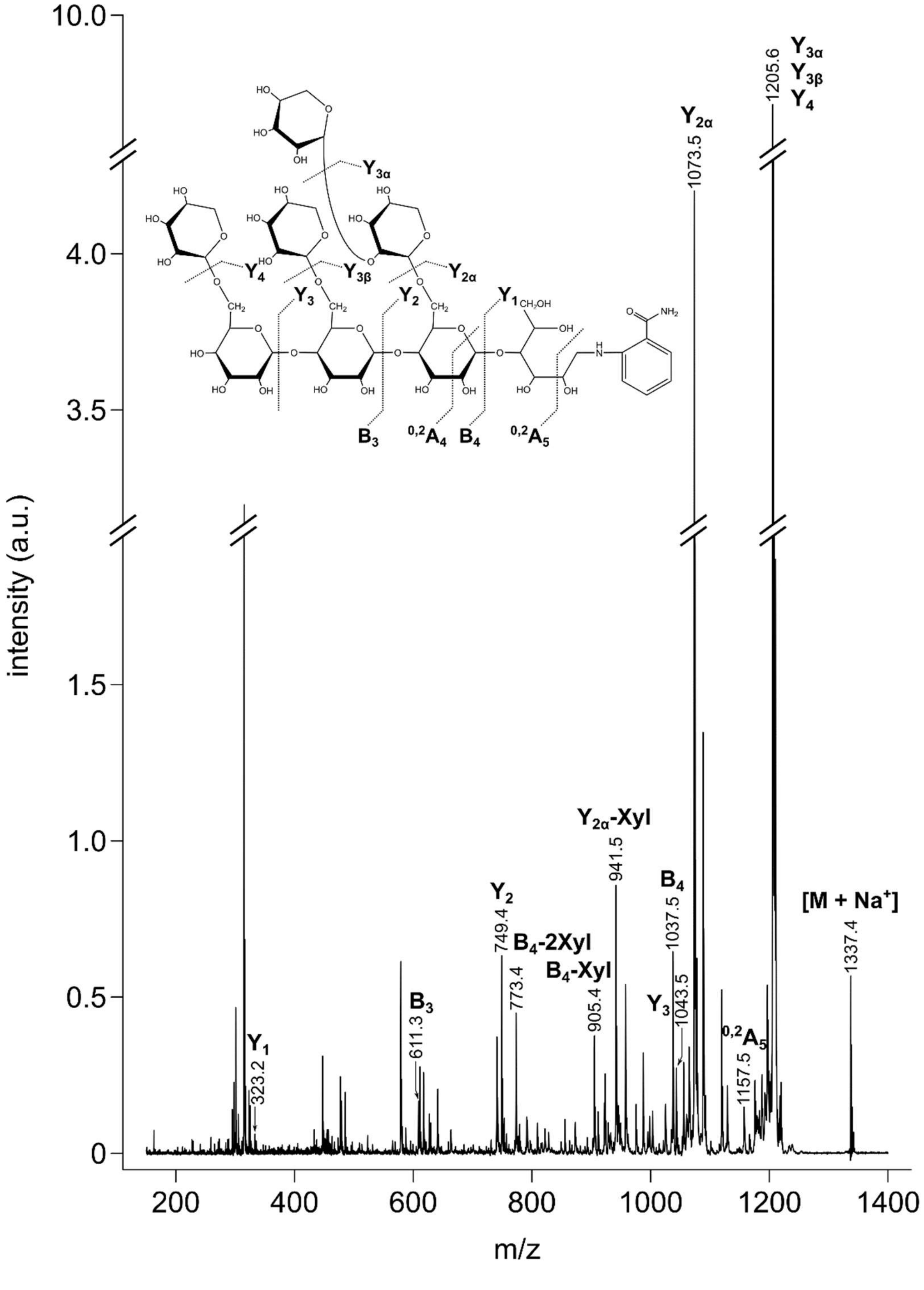

intensity (a.u.)
m/z
10.0
4.0
3.5
1.5
1.0
0.5
0
200
400
600
800
1000
1200
1400
Y3α
Y3β
Y4
1205.6
Y2α
1073.5
Y2α-Xyl
941.5
B4
1037.5
[M + Na+]
1337.4
Y2
749.4
B4-2Xyl
773.4
B4-Xyl
905.4
Y3
1043.5
0,2A5
1157.5
B3
611.3
Y1
323.2
B3
0,2A4
B4
0,2A5

**Fig. S14 Sequential digestion of an unidentified *endo*-XGase product from blueberry skin xyloglucan.** Alkali-extracted hemicellulose from blueberry fruit skin was digested with *Aa*XEG *endo*-XGase before being subjected to ethanol precipitation. A sequential digest was then performed with *Bb*AfcA α1,2-fucosidase (α1,2-Fuc-ase), Fam35 β-galactosidase (β-Gal-ase), *Bo*GH43A, *E. coli* YicI non-reducing-end-specific xyloglucan α1,6-xylosidase (α1,6-Xyl-ase), and *An*GH3 β1,4-glucosidase (β1,4-Glc-ase). After simultaneous treatment with the first three enzymes, and separately after β1,4-Glc-ase digestion, products were isolated using a centrifugal filter (represented by dotted line). For α1,6-Xyl-ase digestions, ethanol precipitation (dotted line) was used to deactivate/remove enzyme.

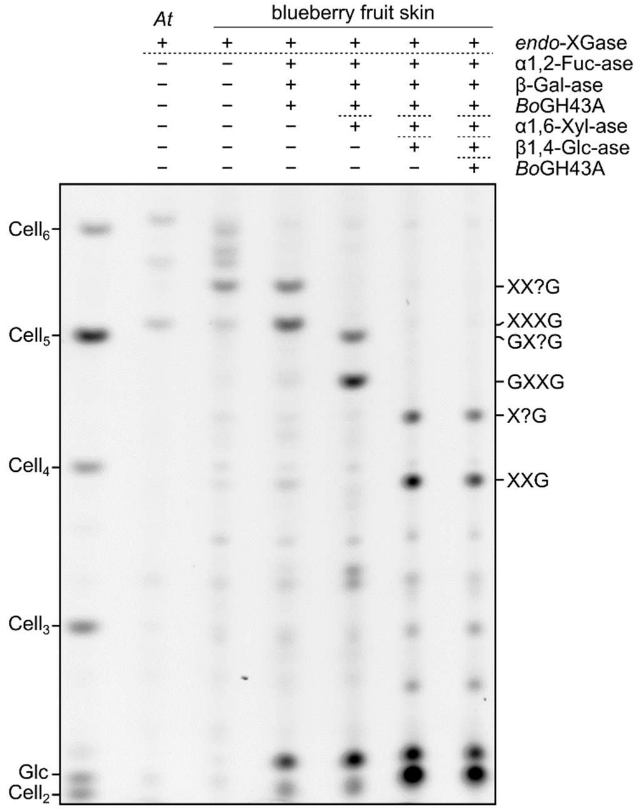

**Fig. S15 Partial characterisation of *endo*-XGase products from *Coffea arabica* leaf xyloglucan.** Alkali-extracted hemicellulose from *C. arabica* leaf was digested with *Aa*XEG *endo*-XGase before being subjected to ethanol precipitation. **a** Left: PACE gel comparing *endo*-XGase products released from xyloglucan of Arabidopsis, tomato, and *C. arabica*. Right: Combinatorial digest of *C. arabica endo*-XGase products with *Bb*AfcA α1,2-fucosidase (α1,2-Fuc-ase), Fam35 β-galactosidase (β-Gal-ase), and/or *Bo*GH43A. **b** MALDI-TOF mass spectrometry analysis of *C. arabica endo*-XGase products. H = unknown hexose; P = unknown pentose.

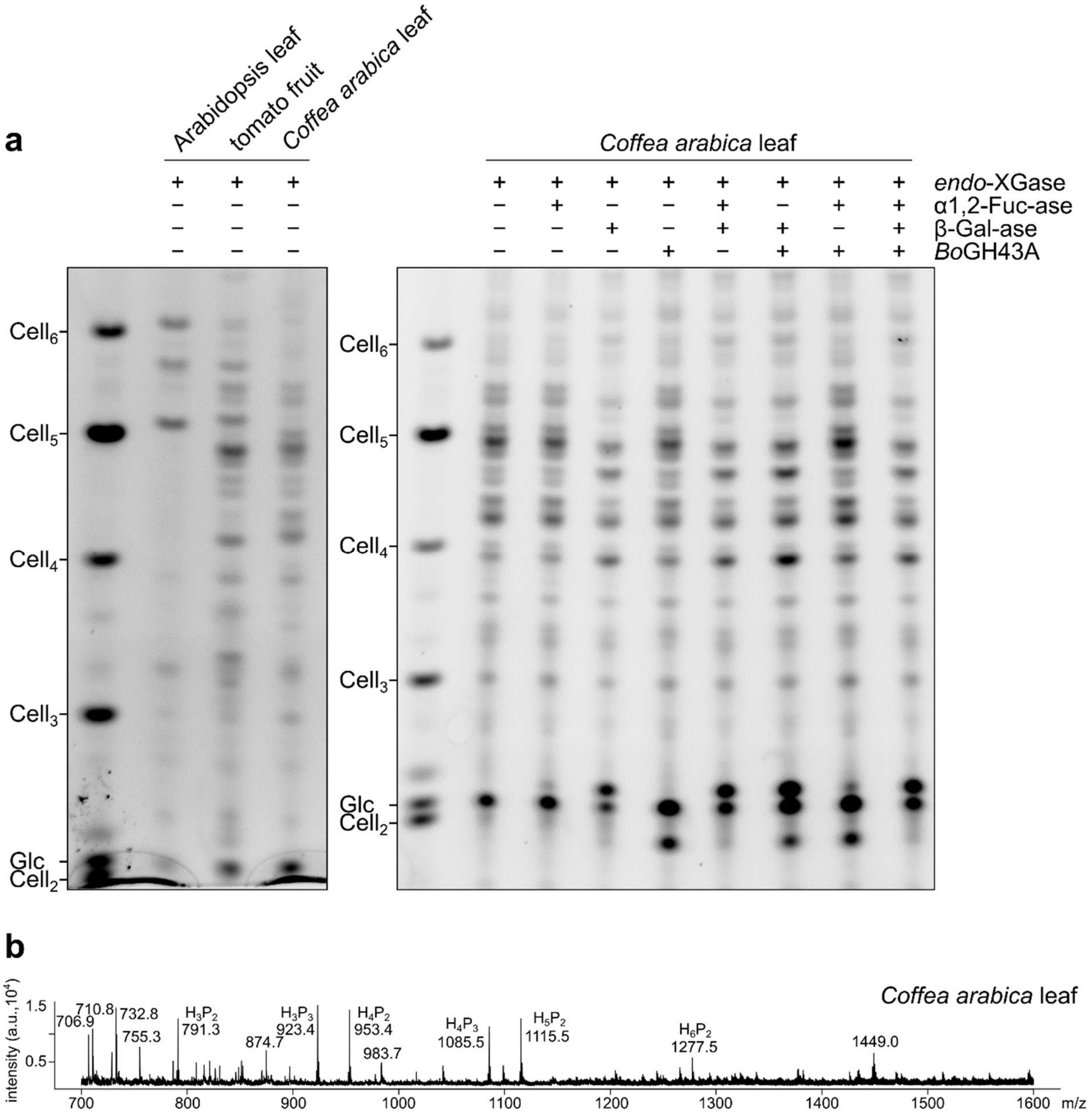

**Fig. S16 No-enzyme controls for *endo*-XGase digestions of plants exhibiting XXGG-/XXG$_n$-type xyloglucan.** Alkali-extracted hemicellulose from transgenic plants was treated with or without *Aa*XEG endo-XGase to confirm that the bands displayed in Fig. 7 result from specific hydrolysis by *endo*-XGase, as opposed to background contamination. Products were subsequently derivatised with a fluorophore and separated by electrophoresis.

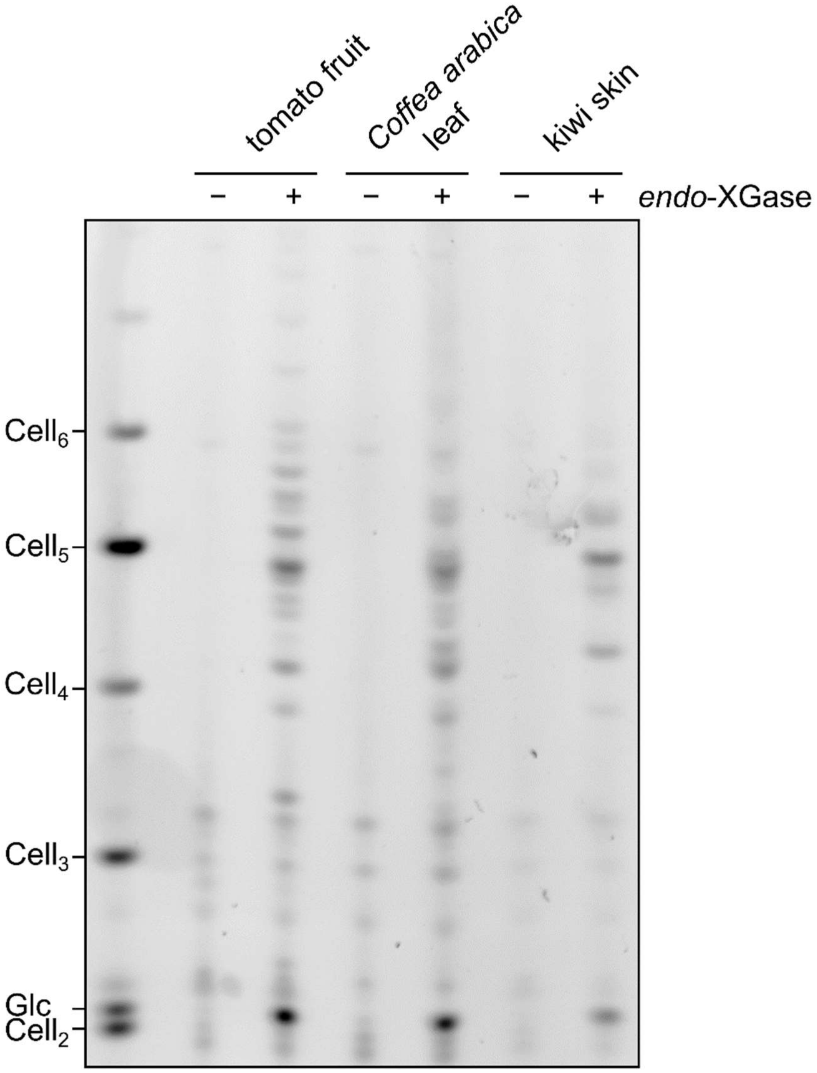

**Fig. S17 Sensitivity of *endo*-XGase products from *Catharanthus roseus* leaf xyloglucan to various *exo*-glycosidases.** Alkali-extracted hemicellulose from *C. roseus* leaf was digested with *Aa*XEG *endo*-XGase before being subjected to ethanol precipitation. The oligosaccharide products underwent a combinatorial digestion with *Bb*AfcA α1,2-fucosidase (α1,2-Fuc-ase), Fam35 β-galactosidase (β-Gal-ase) and *Bo*GH43A. Products were subsequently derivatised with a fluorophore and separated by electrophoresis.

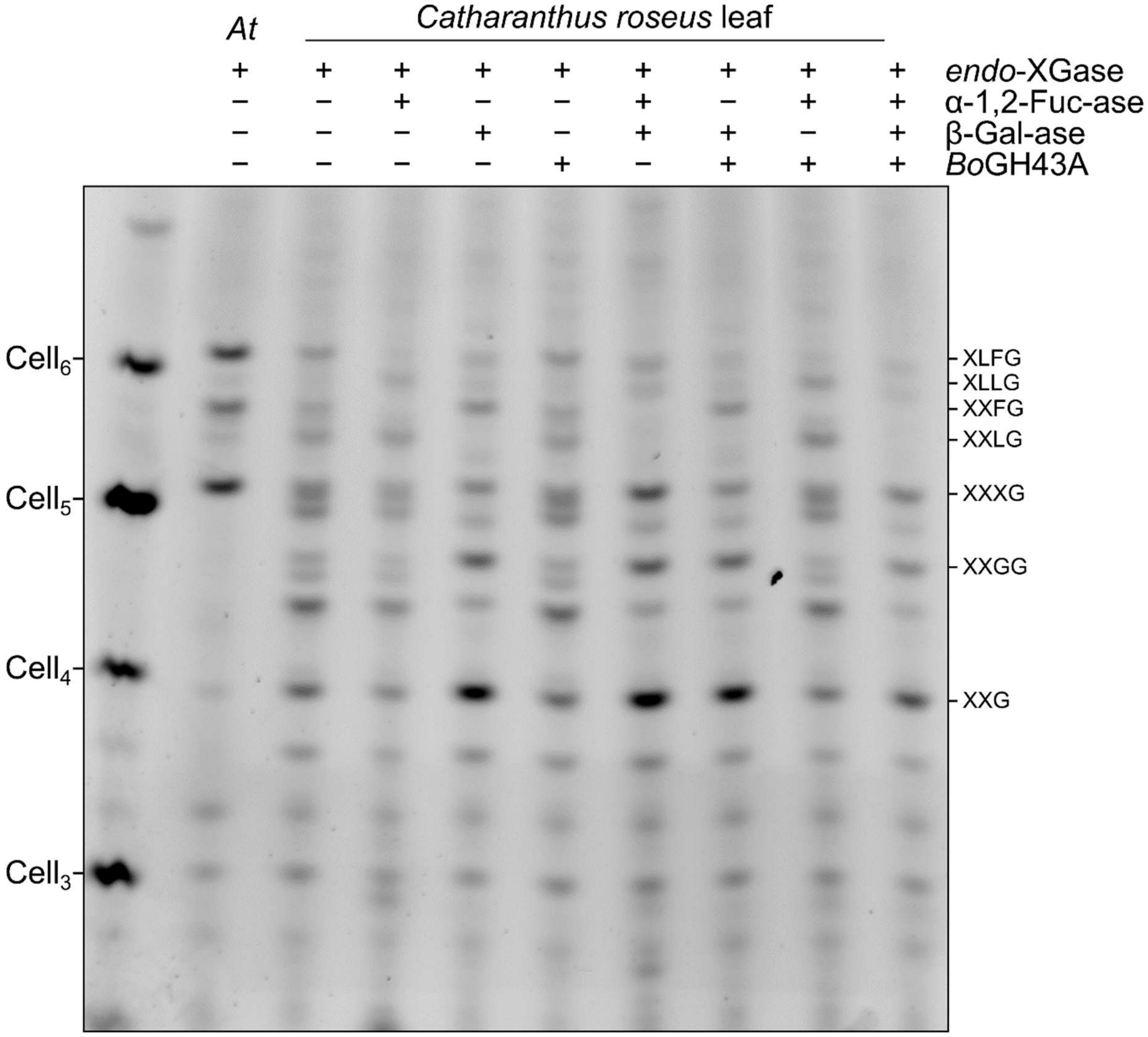

**Fig. S18 Characterisation of *endo*-XGase products from kiwi fruit skin xyloglucan.** Alkali-extracted hemicellulose from kiwi fruit skin was digested with *Aa*XEG *endo*-XGase before being subjected to ethanol precipitation. **a** PACE gel comparing *endo*-XGase products released from xyloglucan of Arabidopsis, tomato, and kiwi. **b** Combinatorial digest of *C. arabica endo*-XGase products with *Bb*AfcA α1,2-fucosidase (α1,2-Fuc-ase), Fam35 β-galactosidase (β-Gal-ase), and/or *Bo*GH43A.

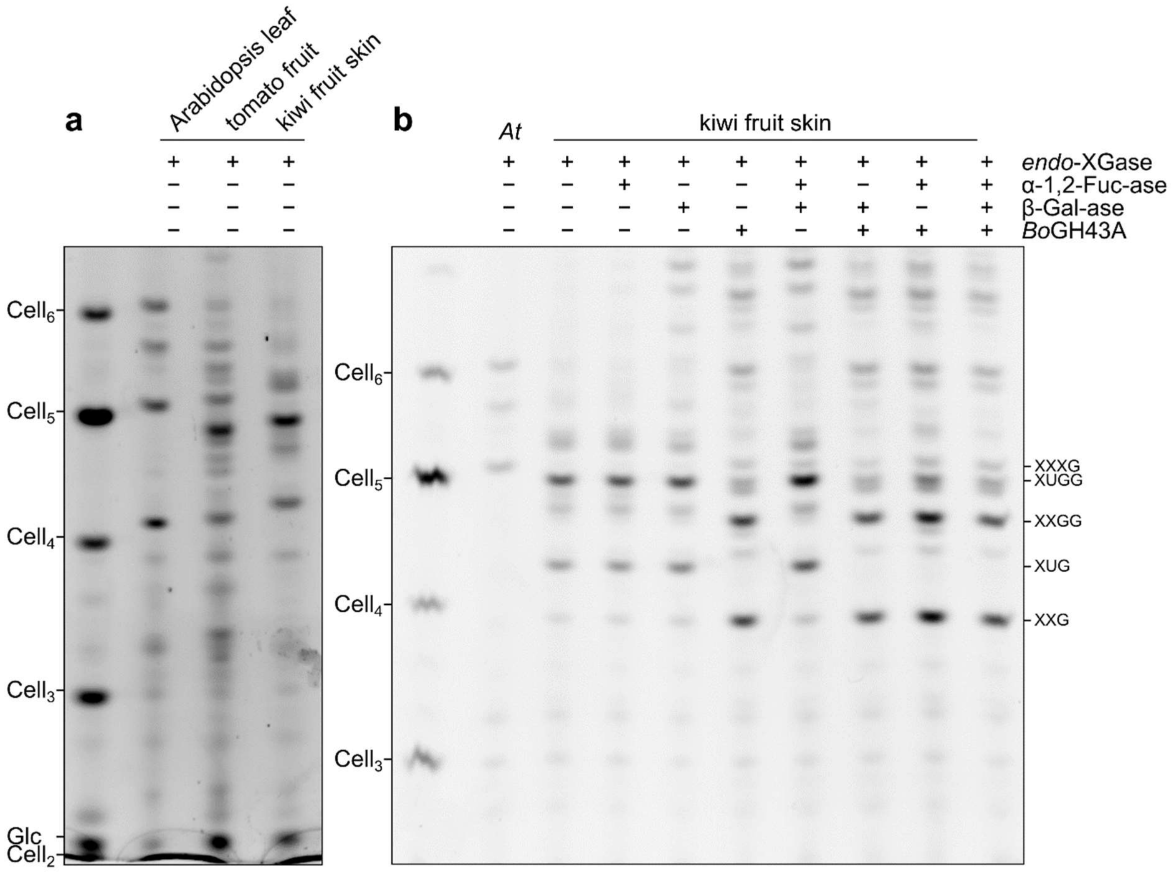


**Table S1** Genomic data sources. Predicted protein sequences (primary transcripts only) were downloaded from the listed genomic data sources. For species marked with an asterisk, only nucleotide sequences were available.

(Table provided separately)

**Table S2** De novo-synthesised coding sequences for Golden Gate assembly. For VmXST1, codons 167–193 (stricken through) were intentionally omitted from the synthesised construct.

### *Vm*XST1

cactctgtggtctcaaATGGCGAATACCTCGTTCCTTCCAACATTTGGCCACCCGTCGACAAATCTCCAAGGCCTAAAAAAGCAGCCGAATAAATCCCCACCCCTTAATT
CCACACTAAAATCATTCCTACACTCCCTTACTCCAACCCGTCGAACGTTAGTCATCATCGTCATCCTTTTCAATGTATTCCTCGTTTTATACCTTACTCGTacccacattc
tctctccatccGAATCCCCTCTAGCATCTCCCGAATTCCTACCCGGAGAATCTAGGATAGTACGAAATATCTCATCGCAAACGTACAGTGGTTCCGGCAGCTTGCAAA
ACAAAGGCGTATTGAATATCTCATCGCAAGCTTACGAGTTACACGACAGCTCGCCGGACTATGACGCACCGGATATCTCATCCCAAACTTATGGCAGATTGCGA
AACAAAGGGGTAGTGGATATCGAATCGCAAAATCACGGCAGCTTGGCGAATAAAGGCGTAGTGAATATCTCATCGGAAAC~~TTACGGCAGCTTGGCGAACAAA~~
~~GAAGTATTGAATATCTCATCGGAAGCGAACAAAGGCATAGTGAATATCTCATCGGAAAG~~TTATGGCAGCTTGGCGAACAAAGAAGTATTGAATATCTCATCGGA
AGCGAACAAAGGCGTAGAGAATATCTCATCGGGACCTTACGGCGGCTTGGCAAACAAAGACGGATTGAATATCTCATCGGAAACTTTCGGTGGCTTGCCGAAC
AAAAGCTGCGATTCCGGTCGGATTTACGTCTATGATCTCCCTCCGATATTGAATTACGATATCAAGAATAATTGCACCCAACTAGAGCCGCCGGACGACAAATGT
GACAAACTTTTGAACGACGGTCTTGGACCGGTGGCGAAAGAGTACGCTGGAATGTTTCCGGAGAGTATTCTCCCTGCGTTCTACTGGACCGATCTTTATTGGGG
CGAGGTGTTGTTTCACAATCGGATGCTTAATCACAGGTGCCGAACACTAGAGCCCGAATCCGCTACGGCGTTCTACATACCTTTTTATGCTGGGCTCGATGTTAG
AAAATACTTGCACAATCACACTGCAAGAGAGCGCGATAGGAATCCTGAGATCCTGACTAAGTGGGTCCAGGAGCAGAAATGGTGGAAGAGATCTAACGGCTCC
GATCACATCATCATGCTTGGAAGGATGACGTGGGATTTCCGACGGTTGGCAGACACTGACGCCGATTGGGGTACCAGATTCCTCAATATGGCCGCGATGCAGA
ATGTCATCCGTGTAACCGTCGAGCGAAACGTGTGGGACAACCTAGAAGTCGCTGTACCCTACCCCACCATATTCCACCCCAGATCAGAATCCGATATCATCCAGT
GGCAGAACTTCCTCAGGAGCCGCCGCCGGAACCACCTCTTCACCTTCGTCGGCGGCGGTCGCTCTAAAATCAAGAACGATTTCCGAGCGGTCCTAAAGAACAAG
TGCCTCAACGAGTCAACCGCGTGCCATCACGTGGACTGCTCCGGGAACAACTGCCTCGACGGGAAAACGGCGGTCATGGCGGCGTTTCTCGACTCCGATTTCTG
TCTCCAGCCGAAGGGGGACGGGTTTACGAGAAGGTCGGTTTTCGATTGCATGTTGGCCGGTTCGGTCCCGGTTTTTTTCTGGAGAGGGACGGCTTACTTGCAAT
ATGAGTTGTACTTCACGCCGAATGAACCGGAGAGTTACTCGGTTTTTATCGACCATAACGACGTGCGCAATGGAACGGATATAAGGAGAGTGTTAGAAGGGTA
TGGGAGAGAGGAAGTGgaaaggatgagagagaaagtaataGAATACATACCGAGATTTGTGTATGCGAAACCGAGTCAAGGGTTGGAGAAAACAAGAGATGCCTTTG
ATATTGCTATTGATGGAGTATTGACCAAATACAAAGCTCACATGGAGAGAGGTAGGGTTGGGAACTGGAATGACATTagttcgtgagaccacgaagtg

### *Cc*XBT1 (Cc07_g06550)

cactctgtggtctcaaATGCTTCCCCTTTCCAATTCCTCGTCACAAGTTAAGGACCGCGGTGTCCATGGGAATCCCAAGCGTACTATTACTGCGTTCGATTTCAAGAAG
GCGCTTAATTTTGTCCGCTCCCATGTTTCTTTCCACCAATTCGCATGGGCCCTCATTCTTTTTCTCTTGGAATTTTGTATCTTCCTAGCCTTCACTGCCACCACTCGTG
CTCCCAGTCCTCCCACAGCCGCCGACCGATTGTCGCCCGTCCGTCGTCCCCAGCTCCAGGGGACGCCGCCGGAACAGGTTAGTCAACAGCCGCTGGTTCCTCCCC
ATCATCCCGATGACCGAGCACAATGCAAGTATGGAAGGGTTTATGTATACGACCTTCCTCCAATTTTCAACAAGAAGCTGCTGGAGAATTGCCAGGACTTAGAC
CCCCGGAGATCCCAGTGCGCGGCGGTGTCCAACGACGGGTTCGGACCAAACGCCACTACGCTCGCCGGCACGGTCCCGCGAGAGCTCGCTCATGCTTGGTACT
GGACCGACCTTTTCGCCGGCGAGATTATCTTTCACGCTCGGATCTCCACCCACAGGTGCAGAACCATGGAGCCAGAATCGGCTGCGGCATTCTACGTGCCCTTTT
ACGCTGGGCTGGCGGTTTCCAAGTATCTGTTCACAAACTACACTGCGAAAGAGCGCGATGCTCCGTGCCAGAATTTGCTCCGGTGGATCAAGGGCCAACCGCAT
TGGAAGAGATCCAACGGTTCTGATCATTTTCTTATGCTGGGGCGTAGTTCGTGGGATTTTAGACGGGCCCGCGATGGAGACTGGGGGACCAGCTTCCTCCTCAT
GCCCCTCATGAGACAAATGTTCCGCTTAACGATAGAAAAGAGCCTTGGCGACCCACTGGAAGTCAGCGTGCCCTATCCCTCCGGATTCCATCCCCGGACCACCTC
CGAAGTTGGGCAGTGGCAGGAATTTGTTCGGAGTCGTAAACGGTCGAGCCTCTTCACGTTTGTTGGCGGAAAACGCGGGTATATCAAGAACGATTTCAGGGCT
TTGTTGCTGGATCAGTGTTACGAGGAGTCGGATTCGTGCAAAGCTGTGGACTGTGCCAAGACCCCATGCTTGGACGGCGCTTCCTCGGTCCTGGACGCCTTTCT
GGACTCGGACTTTTGTCTGCAGCCCAGGGGGGACTCGGTCACCAGGAGGTCCACGTTCGACTGCATGCTGGCCGGCTCCATACCCGTTTTTTTCTGGGAGGGAA
CCGTTGGGGGCCAGTACGAGCTGTACATGTCTGATCAAACCGAGTCCTTTTCGGTTTTCATCCACCGGAATAAAGTGAGGAACGGGACTTCGATAAGGAAAGTG
TTGGAGGGATACAGCAGGGAGGACGTCAAGAGGATGAGGGAGAAAGTGATTGACATGATCCCGAGGATTTCGTACGCTTTTCCAGCTGCGGAGGGAGGATTG
GGCAATCTCAAAGATGCCTTCGACATAGGCGTGGAGGAAGTGCTGAGAAGAATAGTCAAAAATGCCAATCCATACAGGTCGGGAGTCGGGACCCGTATCATGA
ATGAGATCACATTCCCAGAGagttcgtgagaccacgaagtg

## Cc07_g06570

cactctgtggtctcaaATGAAAATGCACCCCTTTGCCAAATTCTCATTATCGTCGTCTGGCACTAGTAGCAGCAGTAGTAGACGCAATTTTGTGGACAAACCCAAGAAC
CGTCGTGCATGGCTTTTCGTTGCTCTCGTCTTTCCTATTCTCATCCTCCTAATGTTCACCAGTGCTCCGAGACAATATCTATCCATCCGTCGTCTGAAGCTACGTTTC
CGAGCTCCGGCTGGTTCTGAGCAGTGCAAGTACGGCACGGTCTATGTATACGACCTTCCTCCATCCTTCAACAAGAAATTGTTAGACAACTGCCGCGCTATAGAT
CCGCAGAATTCACTTTGCGATGCTCTCTCTAACGATGGGTTTGGCCCGAAAGCCACCGATTTCCAGGGAATCATCCCCGAGGATCTTACTCCCGCGTGGTACGCC
ACTAGCATGTTTGCCGGTGAAGTTATTTATCATACCAGGATCTCCAATTACAAGTGCAGAACCTACGATCCAGATTCTGCAGCAGCTTTTTACATACCCTTTTATG
CGGGATTAGCCACTGCCAAGTATCTGTACGCTAATCGCAGTACTGAGAGCGAGAGAGAATCGCAGGGCGAGAGCTTGGTCAAGTGGGTAACTGAGCAACCGT
CTTGGAAGAGATCCAACGGCGCAGATCACTTTATTATGCTGGGCAGAGTATCTGGGGATTTCAGGCCTGGGGATCATCATATGGATGCAAAATCGGGGTCGAG
CTTCGTTCTCATGCCTCCCATCAGACAGACGCTAAGGCTGGCACTTGAACGAAGCCCCTGGGACCGCTACGAGATTGGGGTGCCGTATCCTACGGGATTTCATCC
CAGGTCCAACGCTGAGCTTGAGAAATGGTTGAAATTCGTGAGAACTCGCAACCGGTCCAAGCTCTTCACATTTGTGGGAGGGAAAGAAAAAAGGTTGAAGGAT
GACTTCAGGGGTTTGTTGGTGGACCAGTGCCGCGACGAGTCTGACAGGTGCACAGTTGTGGACTGTTCGGAGACTCCGTGTTCCGATGGAAGCCCGGCAATTC
TTGAACCCTTCCTGGGCTCCAACTTCTGTTTGCAGCCCAGATCAGGTGATTCTTTCACGAGAAGATCCACATTTAATTGCATGCTGGCTGGTTCAATACCGGTTTT
CTTCTGGAAAAGAAGCATCTTTGGTCAGTATGACTGGTTCCTGAGAGATGACCCCGAGCGGTTCTCGGTATTTATAGACGAGAACCAAGTGAGAAACGGGACA
ATTTCTATTAGGAAAGTGTTGGAGGGATATGGCATTGAAGAGATTCAGATGATGAGGGAAAGGGTCACTAATCTGATACCTAGGTTTCTTTATGTTATGCCCGG
TGATGATGAATACGATGATGGTGGTGTGGGGAAGTATACCATGAGAGATGCAGTTGATATAGCTGTTGAAGGGGTGCTGCGACGATTCAAGGAGCTAAGATT
AGTCGAGCGACAAagttcgtgagaccacgaagtg

**Table S3** PCR primers used to amplify DNA parts for Golden Gate assembly.

| Target | | Sequence (5'→3') |
|---|---|---|
| XXT2 promoter | | |
| | Forward | CACTCTGTGGTCTCAGGAGGACTGGTTTGATGTTTTTCATGATATAAACT |
| | Reverse | CACTTCGTGGTCTCACATTCTTCTTTCTTCTTACAAGATTCTCTGTAAAA |
| CESA3 promoter | | |
| | Forward | CACTCTGTGGTCTCAGGAGGGTACGAGAGTTACGAAGCAG |
| | Reverse | CACTTCGTGGTCTCACATTTTGTCACTTAGTTGCTTCCAAC |
| *Sl*XST1 CDS | | |
| | Forward | GTGGTCTCAAATGTTGCCATCTGAAAATTCTTCCCC |
| | Reverse | GTGGTCTCACGAACTTAGTTTTTGTTGCTTGAATCTC |

**Table S4** Exo-glycosidases used in this work.

(Table provided separately)

**Table S5** 1H and 13C NMR assignments for XUXG oligosaccharide.

| Residue | | H1/C1 | H2/C2 | H3/C3 | H4/C4 | H5/C5 | H6/C6 |
|---|---|---|---|---|---|---|---|
| A | α-D-Xyl*p* | 5.141/99.08 | 3.614/81.38 | 3.870/72.69 | 3.652/70.26 | 3.563/3.717/62.07 | |
| B | α-D-Xyl*p* | 4.949/99.51 | 3.542/72.37 | 3.686/73.96 | 3.616/70.39 | 3.571/3.723/62.26 | |
| C | α-D-Xyl*p* | 4.946/99.12 | 3.533/72.28 | 3.713/73.88 | 3.607/70.37 | 3.565/3.732/62.13 | |
| D | β-D-Glc*p* | 4.569/101.23 | 3.45/75.5 | 3.693/74.97 | 3.738/79.97 | 3.818/74.18 | 3.893/3.976/66.9 |
| E | β-D-Glc$p_{nr}$ | 4.56/103.66 | 3.322/73.82 | 3.52/76.29 | 3.539/70.3 | 3.685/75.01 | 3.766/3.950/66.68 |
| F | β-D-Glc*p* | 4.539/103.14 | 3.374/74.39 | 3.666/74.69 | 3.784/79.98 | 3.819/74.71 | 3.908/3.960/66.44 |
| G | β-D-Xyl*p* | 4.53/105.64 | 3.35/74.16 | 3.448/76.45 | 3.644/74.91 | 3.304/3.960/65.91 | |

**Table S6** Kinetic parameters for BoGH43A/B activity on para-nitrophenyl glycosides. Reactions were carried out in 50 mM HEPES, pH 7.5, at 20 °C with 1 µM enzyme. Michaelis-Menten kinetics were collected from initial velocities measured via absorbances at 405 nm. *Parameters extrapolated due to limit of substrate solubility.

| Enzyme | Substrate | $k_{cat}$ ($s^{-1}$) | | $K_M$ (mM) | | $k_{cat}$ / $K_M$ ($M^{-1}s^{-1}$) |
|---|---|---|---|---|---|---|
| *Bo*GH43A | *p*NP-α-Ara*f* | 0.01 | ± 2×10$^{-4}$ | 0.49 | ± 0.04 | 20 |

| | | | | | | |
|---|---|---|---|---|---|---|
| | *p*NP-β-Xyl | 0.1 | ± 4×10$^{-4}$ | 8.9 | ± 0.04 | 10 |
| *Bo*GH43B | *p*NP-α-Ara*f** | 0.01 | ± 2×10$^{-3}$ | 17 | ± 4 | 0.8 |
| | *p*NP-β-Xyl* | 0.02 | ± 2×10$^{-3}$ | 35 | ± 2 | 0.7 |

**Methods S1** Glycosyl hydrolase expression and purification. Plasmids pET-YSBLIC-BoGH43A and pET21a-BoGH43B were transformed into chemically competent E. coli BL21(DE3) (New England Biolabs, Ipswich, MA, USA) according to the manufacturer's protocol and grown on lysogeny broth (LB) agar plates supplemented with 50 mg l−1 kanamycin (BoGH43A) or 100 mg l−1 carbenicillin (BoGH43B) at 37 °C overnight. Colonies were washed off the plate with LB medium and used to inoculate one litre of LB medium (50 mg l−1 kanamycin for BoGH43A or 100 mg l−1 carbenicillin for BoGH43B; baffled flask), which was incubated at 37 °C and 180 rpm until OD600 = 0.7 (BoGH43A) or OD600 = 0.5 (BoGH43B) was reached. Cultures were cooled down, induced with IPTG (0.2 mM for BoGH43A and 0.1 mM for BoGH43B) and shaken (180 rpm) for 18 h at 20 °C (BoGH43A) or 18 °C (BoGH43B). Cultures were harvested by centrifugation at 4000 rpm, 15 °C, for 20 min. Pellets were resuspended in Ni-NTA buffer A (BoGH43A: 50 mM HEPES pH 7, 0.3 M NaCl, 10 mM imidazole; BoGH43B: 50 mM HEPES pH 7, 0.5 M NaCl, 30 mM imidazole) supplemented with 2.5 mM MgCl2, Benzonase (5 µl of the solution provided by the supplier; Merck, Darmstadt, Germany) and 1 mg ml−1 lysozyme. Following incubation for 30 min rolling at 4 °C, lysis was completed by sonication (20 cycles, 10 s pulse, 30 s off) on ice. The supernatant was cleared by centrifugation (12,000 rpm, 4 °C , 40 min) and loaded onto a Ni-NTA gravity column (column volume, CV: 2 ml) equilibrated in Ni-NTA buffer A. The column was washed with 30 CV Ni-NTA wash buffer (BoGH43A: 50 mM HEPES pH 7, 0.3 M NaCl, 20 mM imidazole; BoGH43B: 50 mM HEPES pH 7, 0.5 M NaCl, 30 mM imidazole) and protein eluted with 3 CV Ni-NTA elution buffer (BoGH43A: 50 mM HEPES pH 7, 0.3 M NaCl, 500 mM imidazole; BoGH43B: 50 mM HEPES pH 7, 0.5 M NaCl, 500 mM imidazole). The protein solution was concentrated in an Amicon centrifugal filter (30 kDa cut-off; Merck), buffer exchanged to size-exclusion (SEC) buffer (BoGH43A: 25 mM HEPES pH 7, 0.1 M NaCl, 1 mM DTT; BoGH43B: 10 mM HEPES pH 7, 0.25 M NaCl) using a PD-10 column (GE) and concentrated in an Amicon centrifugal filter (30 kDa cut-off; Merck) to 1 ml. The protein solution was loaded onto a HiLoad 16/60 Superdex 200 prep grade (GE) equilibrated in SEC buffer and after a 40 ml void volume 1 ml fractions were collected. Fractions containing BoGH43A or BoGH43B were pooled and concentrated in a 30 kDa Amicon centrifugal filter (Merck) to 15 mg ml−1 (BoGH43A) or 10 mg ml−1 (BoGH43B). For storage, glycerol was added (25% final concentration), protein was frozen in liquid nitrogen and stored at −80 °C.

**Methods S2** Oligosaccharide purification. To purify the XUXG oligosaccharide, deacetylated hemicellulose was extracted from rosette leaf AIR of six-week-old CcXBT1-expressing plants (as described above), applying 100 mg AIR to the PD-10 column. To the 3.5 ml eluate, 50 µl 14 µM AaXEG, 20 µl 9.5 µM BbAfcA α1,2-fucosidase, and 20 µl 25 µM Fam35 β-galactosidase were added before incubating at 37 °C for 18 h. The digest was then dried and resuspended in a total of 100 µl 50 mM ammonium acetate, pH 6.0. This sample was applied to a 500 ml Bio-Gel P-2 (BioRad, Hercules, CA, USA) gravity column equilibrated with the same ammonium acetate buffer. Two-millilitre fractions were collected at a flow rate of approximately 0.1 ml min−1. Fractions were analysed by drying a 25 µl aliquot, labelling in 5 µl fluorescent labelling reagent and visualising by PACE (as described above). The XUXG-containing fractions were pooled, further purified using a PD MiniTrap G-10 column (Cytiva, Marlborough, MA, USA) equilibrated to the same buffer, and dried. A similar method was used to enrich the blueberry XyGO of interest, applying 200 mg blueberry skin AIR to each of two PD-10 columns. To each 3.5 ml eluate, 75 µl 14 µM AaXEG was added, and the samples were incubated at 37 °C for 18 h. For precipitation, 6.5 ml of ethanol was added to both aliquots, followed by incubation, centrifugation, and drying of the supernatant as described above. Both samples were then combined in a total volume of 200 µl ammonium acetate buffer; subsequently, 30 µl 9.5 µM BbAfcA, 30 µl 25 µM Fam35 β-galactosidase, and 15 µl 25 µM BoGH43A were added. The reaction was incubated at 37 °C overnight. Ethanol was then added to a final concentration of 70 % before another round of precipitation, centrifugation, and drying of the supernatant. The sample was then resuspended in 1 ml ammonium acetate buffer; size exclusion chromatography then proceeded as above.
